# MIRO2 regulates prostate cancer cell growth via GCN1-dependent stress signaling

**DOI:** 10.1101/2021.05.20.444992

**Authors:** Madison Furnish, Dillon P. Boulton, Victoria Genther, Mitchell Lee Ellinwood, Lina Romero, M. Scott Lucia, Scott D. Cramer, M. Cecilia Caino

## Abstract

There is a continued need to identify novel therapeutic targets to prevent the mortality associated with prostate cancer. In this context, we discovered a novel mitochondrial signaling pathway that controls androgen-independent and androgen-sensitive prostate cancer cell growth. Mitochondrial Rho GTPase 2 (*MIRO2*) mRNA was upregulated in prostate cancer compared to localized tumors, and higher *MIRO2* levels were correlated with poor patient survival. Using human cell lines that represent AR-independent or androgen-sensitive prostate cancer, we show that MIRO2 depletion impaired cell growth, colony formation and tumor growth in mice. Network analysis of MIRO2’s binding partners identified metabolism, cell cycle, and cellular responses to extracellular stimuli amongst the top over-represented pathways. The top hit on our screen was General Control Non-derepressible 1 (GCN1). *GCN1* was overexpressed in prostate cancer and MIRO2-GCN1 interacted in prostate cancer cell lines and in primary prostate cancer cells. Our results showed that MIRO2 is necessary for efficient GCN1-mediated GCN2 kinase activation and signaling, triggering translation of the transcription factor ATF4. Importantly, MIRO2 controlled ATF4 levels and transcriptional activity both in amino acid replete and depleted conditions. Furthermore, MIRO2’s effect on regulating prostate cancer cell growth was partially mediated by ATF4. Finally, activation of GCN2 and ATF4 expression were correlated with MIRO2 expression in prostate cancer xenografts. Overall, we propose a new mechanism driving prostate cancer growth of both AR-independent and androgen-sensitive tumors.

## INTRODUCTION

Patients diagnosed with metastatic prostate cancer face a 5-year survival rate of 30.2% (SEER Program, NCI). Hormone therapy targeting the androgen axis (therapeutic castration) is the mainstay therapy in prostate cancer (1). Because many tumors become resistant to castration but continue to be driven by androgen signaling, novel drugs that block androgen production or target the androgen receptor (AR) are used in the clinic (2–4). Unfortunately, resistance to second generation AR-targeted therapies and progression to androgen-independent disease ultimately leads to prostate cancer-related death (3–7). For these patients, there are no effective treatments, highlighting the continued need to identify novel therapeutic targets to prevent the mortality associated with prostate cancer.

In this context, mitochondria play a central and multifunctional role in malignant tumor progression (8). In particular, mitochondrial signaling pathways have emerged as important nodes for tumorigenesis and metastatic dissemination (9–12). As a result, mitochondria-targeting strategies are being pursued in the clinic and there is a strong interest in identifying new mitochondrial pathways in cancer (8, 10). In this study, we focus on one novel mitochondrial pathway centered on mitochondrial Rho GTPase 2 (MIRO2, also known as Ras Homolog Family Member T2, RHOT2), a small GTPase of the Ras-superfamily. In previous studies, we showed that *MIRO2* mRNA is up-regulated in primary cancer *versus* normal tissues, across multiple tumor types (13). MIRO2 had been previously studied in the context of non-tumorigenic cells, where it coordinates microtubule- and actin-based mitochondrial movement (14). However, we showed that cancer cells utilize MIRO2’s canonical function on mitochondrial trafficking exclusively under conditions of cellular stress that exacerbate mitochondrial dynamics (13, 15). In a recent study, Altieri and colleagues showed that Myc transcriptionally regulates *MIRO2*, which in turn modulates mitochondrial dynamics and tumor cell invasion (16). Despite this evidence for a role of MIRO2 in cancer, we lack a comprehensive understanding of the function and regulation of MIRO2 in cancer cell biology.

The goals of this study were to examine the role of MIRO2 in driving tumor cell-intrinsic phenotypes in prostate cancer and to characterize the molecular pathway utilized by MIRO2 in this context. In prostate, *MIRO2* mRNA is upregulated in patients with recurred or progressed disease, and higher *MIRO2* levels correlate with poor patient survival. MIRO2 depletion in prostate cancer cell lines impaired cell growth, colony formation, and tumor growth in mice. Network analyses of MIRO2 co-precipitating proteins identified metabolism, cell cycle and cellular responses to extracellular stimuli amongst the top over-represented pathways. We characterized the role of the top hit on our screening, General Control Non-derepressible (GCN1) and found that GCN1 depletion mimicked MIRO2 depletion. Furthermore, MIRO2 controlled activation of GCN1 signaling *in vitro* and *in vivo*. Overall, we propose a new mechanism driving prostate cancer growth of both AR-independent and androgen-sensitive tumors.

## MATERIALS AND METHODS

**Supplementary information Methods** include detailed sections on Cell culture, Antibodies and reagents, Plasmids and transfections, Gene silencing, Western blotting, Endogenous co-IP, FLAG IP, Proteomics ID, mRNA quantitation, Analyses of expression and cancer cell dependency from public databases, Cell growth assays, Cell Cycle analyses, Proximity ligation assays, Immunofluorescence and cortical mitochondria quantitation, and Immunohistochemistry.

### Primary culture of human prostatic epithelial cells

Monolayer cultures of human prostatic epithelial cells were conducted as previously described (17) with minor modifications. Samples were obtained through an honest broker through the Pathology Shared Resource of the University of Colorado Cancer Center. The authors did not have access to patient identifiers and the study was considered exempt from human subjects research by the Colorado Institutional Review Board. Briefly, areas of benign and tumor glands in a fresh prostatic cross section were identified by toluidine blue staining. Samples were obtained within one hour of surgical removal. The remaining areas where samples were removed were included in routine diagnostic blocks and H&E sections were used to assess glandular morphology and assign Gleason grades. Each patient sample was given a unique identifier (i.e., UCD1 or UCD2). Cultures grown from areas of greater than 90% tumor surrounding a location were designated cancer and were assigned CA to their name (i.e., UCD1-CA or UCD2-CA). Cultures grown from areas surrounded by 100% benign glands were designated benign and assigned B to their name (i.e., UCD1-B or UCD2-B). The fresh samples were processed exactly as described previously (17). Primary explants were grown on collagen coated dishes in serum-free KSFM medium with supplements (Gibco). After primary outgrowth, the cells were frozen in aliquots for later use. Secondary cultures were grown on collagen-coated plates in MCDB105 (Gibco) plus supplements. All cultures were used at passage 2 or 3.

### Animal studies

Size of experimental groups were calculated to detect a minimal difference of 40% between control and experimental groups, and 30% variability around the mean. Under these conditions, at 9 animals per group with expected effect size of 0.4 between the two compared groups at the two-sided 0.05 significance level, we have 80% power to detect differences. Experimenters were blinded until the end of the study, with one person preparing the cell suspensions, and a second injecting animals and measuring tumors). Groups of 8-week-old outbred male immunocompromised athymic mice (Foxn1^nu^/Foxn1^nu^, Jackson Laboratory, strain##007850 J:NU) (9 mice per group) were injected s.c. with 1×10^6^ PC3 cells stably transfected with control empty vector or two independent MIRO2-directed shRNA sequences (M2 a and b), and superficial tumor growth was quantified with a caliper. At the end of the experiment, animals were euthanized, and the xenografts were dissected and processed for immunohistochemistry (IHC).

### Statistics

Experiments were carried out in triplicates and data are expressed as mean ± SEM of multiple independent experiments (at least 3 independent experiments, n=3). For descriptive data analysis, means, S.D. and medians were calculated, and distributions of data were examined to ascertain whether normal theory methods were appropriate. Student’s t-test or Wilcoxon rank sum test was used for two-group comparative analyses. For multiple-group comparisons, ANOVA or Kruskal–Wallis test with post-hoc Bonferroni’s procedure were applied. Variance similarity between groups was tested with Fisher (two groups) or Bartlett’s (multiple groups) tests. All statistical analyses were performed using GraphPad software package (Prism 7.0) for Windows. A *p* value of <0.05 was considered as statistically significant.

### Study approvals

Studies involving vertebrate animals (rodents) were carried out in accordance with the Guide for the Care and Use of Laboratory Animals (National Academies Press, 2011). Protocols were approved by the Institutional Animal Care and Use Committee of the University of Colorado (IACUC protocol #581). Studies using human primary prostate epithelial cells were considered exempt from human subjects research by the Colorado Institutional Review Board.

## RESULTS

### *MIRO2* alterations in prostate cancer

To examine the status of *MIRO2* in cancer, we searched the TCGA PanCancer Atlas database at cBioPortal (18, 19) for *MIRO2* expression, copy number alterations and mutations. We also investigated the status of the closely related *MIRO1*, as studies in non-tumorigenic cells have shown functional redundancy between MIRO1 and MIRO2 (14, 20, 21). We found that *MIRO2* mRNA is expressed at variable levels across tumor types, with prostate adenocarcinoma (PRAD) showing one of the highest medians (Fig. 1a). Interestingly, *MIRO2* is expressed at higher levels than *MIRO1* in the same tumor types (Fig. 1a). In the TCGA PRAD cohort, *MIRO1* and *MIRO2* expression are oppositely regulated in cancer *versus* normal adjacent tissue (Fig. 1b). Likewise, higher *MIRO2* expression in prostate cancer compared to normal prostate is seen in other patient cohorts, including the Grasso and Lapointe datasets (Fig. S1a). Furthermore, higher *MIRO2* expression is correlated with poor patient survival in the TCGA cohort, while there is no correlation with *MIRO1* expression in the same patient cohort (Fig. 1c). A second patient cohort showed a trend for higher levels of *MIRO2* association with poor patient survival (Fig. S1d). Recurred or progressive disease, as well as metastatic disease, were associated with higher *MIRO2* expression as well (Fig. 1d and Fig. S1b). On the other hand, there is no relationship between expression of *MIRO2* and Gleason grade of the tumors (Fig. 1e and Fig. S1c). At the genomic level, *MIRO2* is commonly altered in prostate, breast and pancreas tumors (Fig. S1e). Mutations in *MIRO2* are randomly distributed, with a small number of more frequent sites associated with functional domains (Fig. S1f). Overall, *MIRO2* is expressed across disparate tumor types and upregulation in prostate cancer was associated with poor prognosis and metastatic disease.

**Figure 1.**
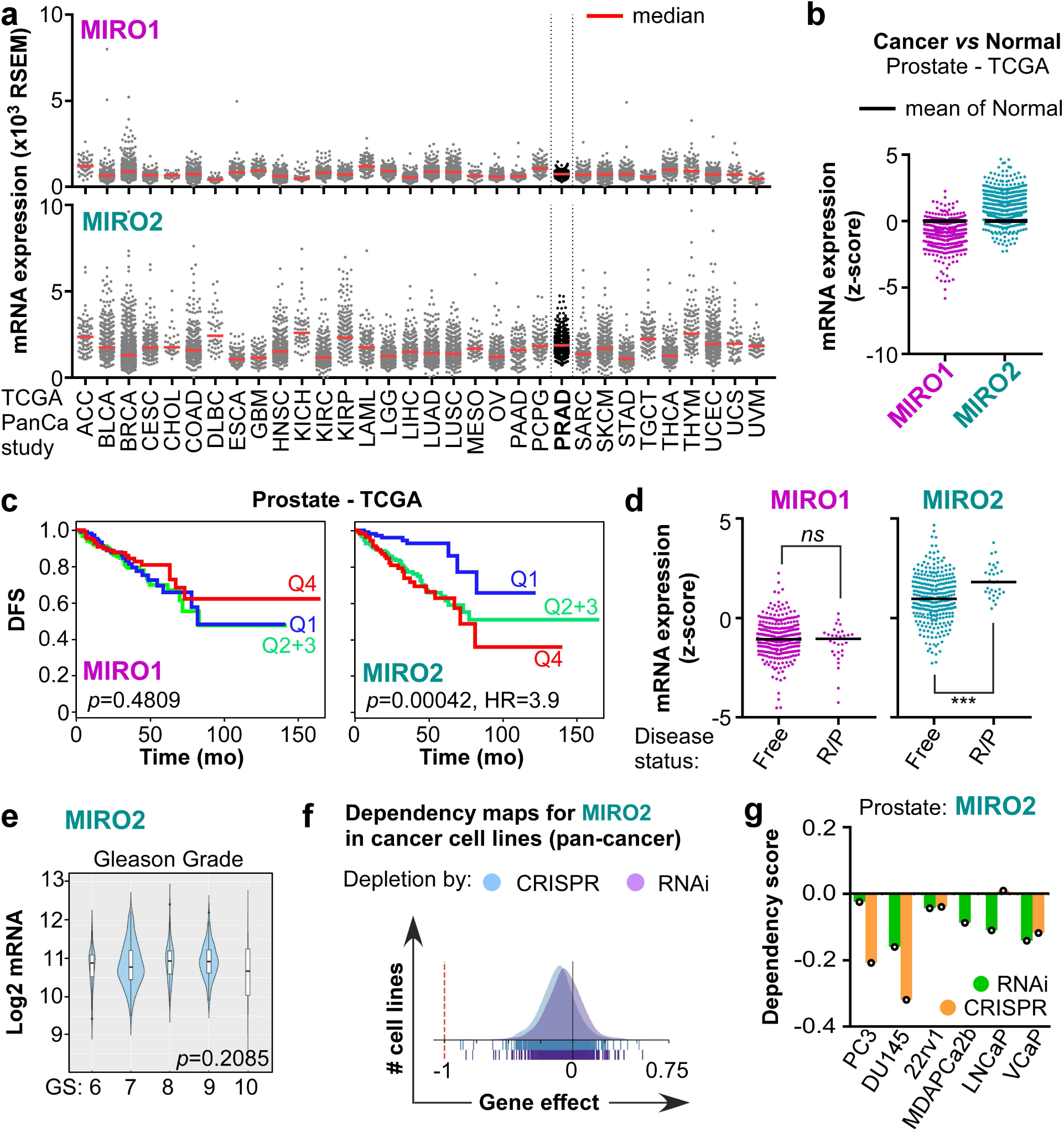
MIRO1/2 alterations in cancer. **a**, The TCGA database was interrogated for *MIRO1/2* mRNA expression across tumor types. **b**, Relative expression of *MIRO1*/*2* mRNA in cancer *versus* normal adjacent tissues on the prostate TCGA PanCancer study. **c**, Kaplan-Meier analyses based on *MIRO1/2* mRNA expression in human primary tumor samples. Datasets were split at quartiles, where Q4>Q3>Q2>Q1. Q1 vs Q4 curves were compared with a Mantel-Cox test. Hazard ratio (HR) was calculated with a Cox proportional hazards regression model. DFS, disease-free survival. **d**, *MIRO1/2* expression according to disease free status. R/P, recurred/progressed. Ns, not significant. ***, p<0.0001 by t-test with Welch’s correction. **e**, Violin plots depicting *MIRO2* expression among the Prostate TCGA dataset according to Gleason grade were generated using CANCERTOOL. GS, Gleason score. Groups were compared by ANOVA. **f**, The DepMap portal was searched for genetic cancer dependency on *MIRO2*. Gene effect scores are derived from DEMETER2 or CERES, with lower scores meaning a cell line is more likely to be dependent in the gene. A score of 0 represents non-essential genes, while −1 corresponds to the median of all common essential genes. **g**, Dependency scores in prostate cancer cell lines subject to RNAi or CRISPR-mediated depletion of MIRO2.

As a first way to examine genetic dependency of cancer cells on *MIRO2*, we turned to the Cancer Dependency Map project from the Broad Institute. Depletion of *MIRO2* in a large panel of pan-cancer cell lines by RNAi or CRISPR/Cas9 gene editing reduced cancer cell growth in the majority of cell lines tested (Fig. 1f). In a prostate cancer cell line panel, *MIRO2* depletion reduced cell growth (score < 0) in all cell lines tested (Fig. 1g). In summary, *MIRO2* is overexpressed in prostate cancer and prostate cancer cells are uniformly dependent in *MIRO2* for cell growth.

### MIRO2 controls prostate cancer cell growth

In prostate cancer cell lines that do not express AR (PC3 and DU145), MIRO2 depletion impaired anchorage-dependent (Fig. 2a,b) and anchorage-independent cell growth (Fig. 2c). Cell viability assays showed significant impairment of growth kinetics in MIRO2-depleted AR-independent prostate cancer cells (Fig. 2d). Furthermore, cell viability was impaired in the androgen-sensitive cell lines(22)(23) C4-2 and 22rv1 (Fig. 2e,f). In contrast, MIRO1 depletion had no effect on anchorage-dependent growth (Fig. S2a,b) or cell viability (Fig. S2c-e), indicating non-redundant roles for MIRO1 and MIRO2 in prostate cancer cell biology.

**Figure 2.**
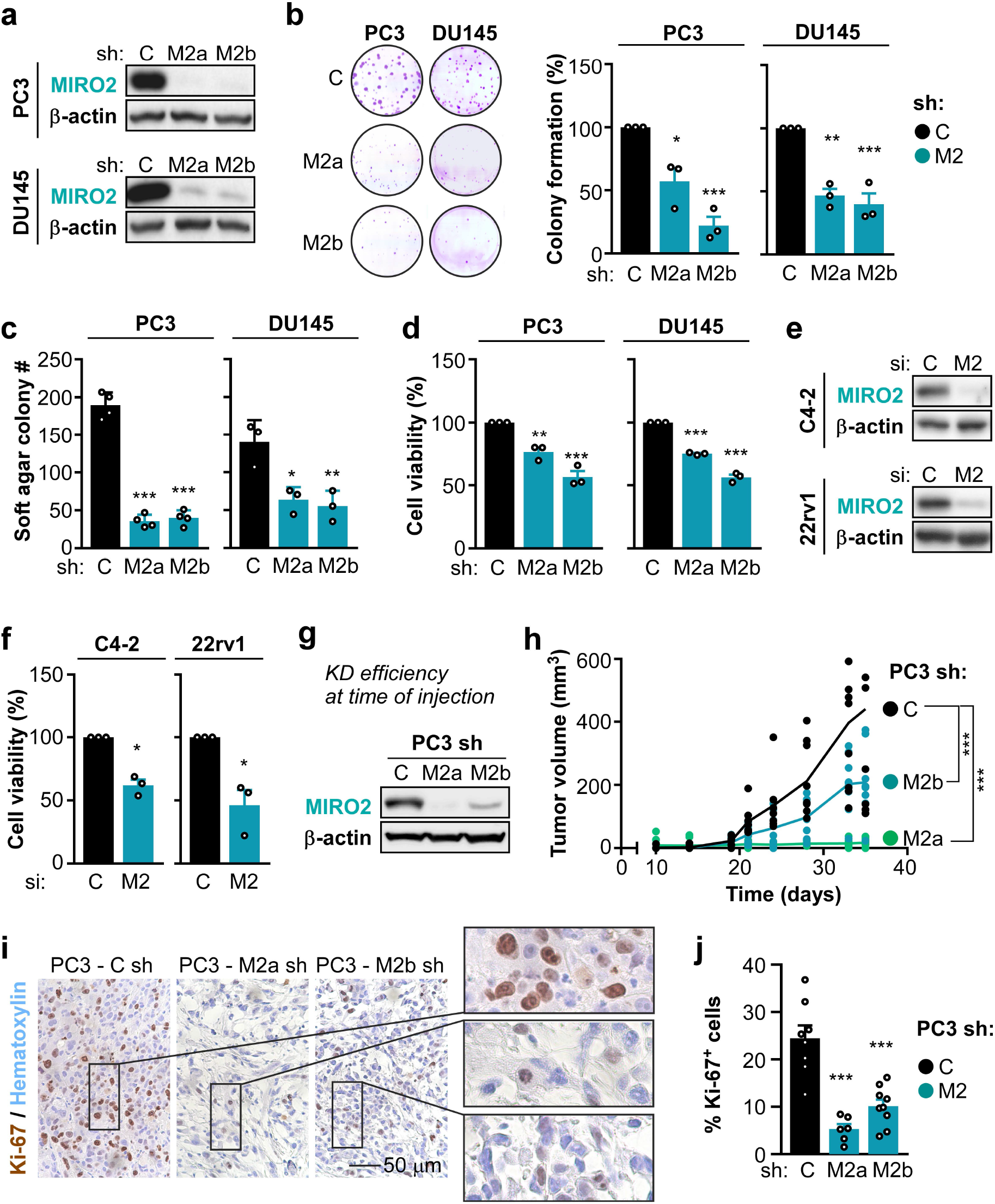
MIRO2 depletion impairs tumor cell intrinsic phenotypes. Stable knockdown of MIRO2 was achieved by shRNA in PC3 and DU145 cells. C, control sh; M2a/b, MIRO2-targeting shRNAs (two independent sequences, a and b). **a**, Representative blots showing the efficiency of knockdown. **b**, Anchorage-dependent growth at 14 days post-plating. *Left*, Representative scans of stained colonies. *Right*, quantitation of colony # per well, relative to control (C) and represented as mean ± SEM (n=3). *, *p*=0.0112; ***, *p*<0.001 by One-Way ANOVA and Dunnett’s post-test for multiple comparisons. **c**, Anchorage-independent growth in soft agar at 14 days post-plating. Colony # per well was quantitated and represented as mean ± SEM (n=3). ***, *p*<0.001 by One-Way ANOVA and Dunnett’s post-test for multiple comparisons. **d**, Cell viability was determined by a MT-Glo assay and relativized to the control (Csh). Data is represented as mean ± SEM (n=3). **, *p*<0.01; ***, *p*<0.001 by One-Way ANOVA and Dunnett’s post-test for multiple comparisons. **e-f**, Cells were transiently transfected with control (C) or MIRO2 (M2)-targeting siRNA. **e**, Representative WB showing the level of depletion of MIRO2. **f**, Cell viability was determined 6 days later by a MT-Glo assay and relativized to the control (Csi). *Left*, data is represented as mean ± SEM (n=3). *, *p*<0.05 by unpaired t-test with Welch’s correction. **g-h**, Cells were injected *s.c.* into the flanks of male nude mice and tumor growth was followed by caliper measurements. **g**, Immediately after injection, cells were pelleted and analyzed for efficiency of knockdown by Western blot. **h**, Data are represented as individual tumors with the line connecting the mean of each group (n=9 for control or M2b, n=6 for M2a). Only 6/9 injected animals developed tumor in the M2a group. ***, *p*<0.001 by 2-Way ANOVA. **i**, Tumors from (h) were processed for IHC at the endpoint of the experiment and assayed for proliferation by staining with Ki-67 antibody and hematoxylin. A representative field (magnification 40X) is shown. **j**, Data are represented as mean ± SEM (n=9 for control or M2b, n=6 for M2a). ***, *p*<0.001 by One-Way ANOVA and Dunnett’s post-test for multiple comparisons.

Next, we evaluated the kinetics of tumor growth *in vivo*. We found that MIRO2 depletion impaired tumor growth in mice (Fig. 2g,h). Furthermore, the proliferative index of MIRO2-depleted tumors was significantly lower compared to control (Fig. 2i,j). This suggests that MIRO2 supports tumor growth by stimulating cell cycle progression. To confirm this *in vitro*, we analyzed cell cycle profiles in prostate cancer cell lines growing in culture conditions. We found that MIRO2 depletion led to an increase in cells in the G2/M phase of the cell cycle (Fig. S2f,g). In sum, MIRO2 controls several tumor cell-intrinsic phenotypes that support prostate cancer cell growth *in vitro* and *in vivo*.

### Novel effectors of MIRO2 in prostate cancer

A key question stemming from our studies was what are the mechanisms by which MIRO2 supports tumor cell growth in prostate cancer? As MIRO2 has been linked to mitochondrial subcellular localization at the cortical cytoskeleton in response to therapy and oncogene-induced stress (13, 15, 16), we first analyzed the distribution of mitochondria in control and MIRO2-depleted cells growing in normal conditions (e.g. non-stressed). Using a panel of prostate cancer, glioblastoma, breast and leukemia cells, we found no differences in cortical mitochondrial localization between control and MIRO2-depleted cells (Fig. S3a,b). Thus, cell growth defects are not associated with alterations in MIRO2-dependent mitochondrial localization under normal growth conditions.

Recent studies in non-tumorigenic cells suggest that MIRO2 might have overlapping functions with MIRO1 in regulating mito-ER contacts, mitophagy and mitochondrial shape (14, 21, 24–26). As contextual signals for neurons, hepatocytes or fibroblasts likely do not overlap with those for epithelial cancer cells, we decided to take an unbiased approach to learn the molecular function of MIRO2 in cancer. MIRO2 is a member of the Ras-superfamily of small GTPases (27). Small GTPases are molecular switches that bind effectors, which in turn activate signaling cascades and gene expression programs (28–30). Thus, we reasoned that identifying MIRO2’s protein binding partners would shed light on the molecular function of MIRO2 in prostate cancer.

To this end, we identified the proteins co-precipitating with FLAG-MIRO2 from PC3 cells (Tables S1, S2). Network analysis of MIRO2 binding partners identified metabolism, cell cycle and cellular responses to extracellular stimuli amongst the top over-represented pathways (Fig. 3a and Table S3). The top 50 co-IP protein candidates based on spectral counts are presented in Table S1. These include metabolism-related enzymes, protein kinases and protein kinases regulators (GCN1, mTOR, SYFA, TECR, CMC1, SFXN3, QCR2, NDUS1, CMC2, FADS2, SQOR, ACSL3, M2OM); ubiquitin ligases/deubiquitinases and proteasome-related proteins (RN213, HUWE1, UBR4, USP9X, ECM29, BIRC6, FAF2); chaperones and co-chaperones (BAG6, MDN1, DNJA1); organelle structure, post-translational modification and trafficking proteins (BIG1, RPN2, MON2, EHD4, GET4, EHD1); and organelle import/export proteins (XPO1, XPO2, NU205, IPO5, IPO7). We validated candidate proteins ranging in spectral counts between 20-200 in FLAG-MIRO2 co-IP and found that 6/6 were confirmed to co-IP with FLAG-MIRO2 using endogenous antibodies for the protein candidates (Fig. S3c).

**Figure 3.**
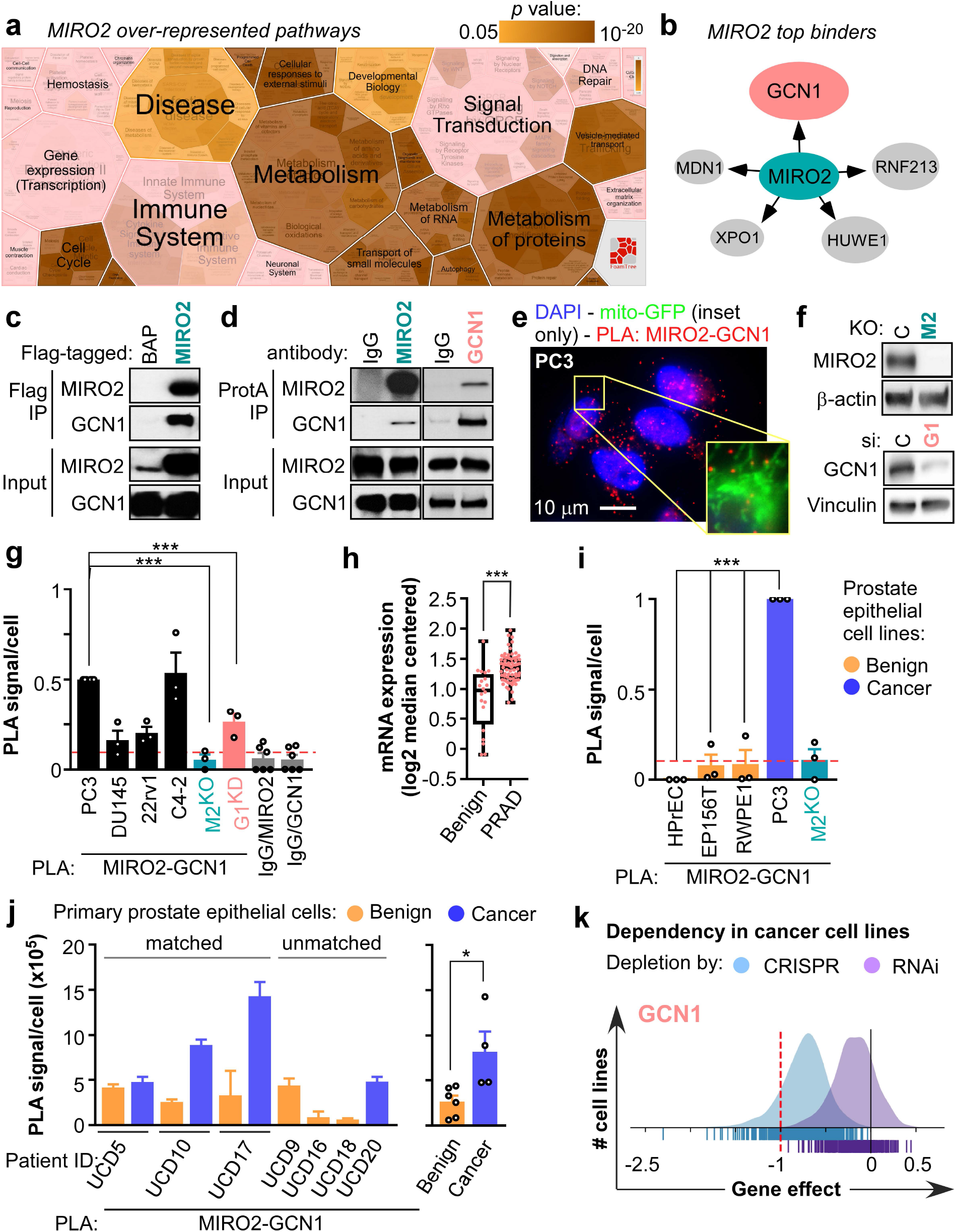
Identification of novel effectors of MIRO2 in prostate cancer. **a**, MIRO2-Flag was immunoprecipitated (IP) from PC3 cells and protein binders were tryptically digested and identified by shotgun proteomics. A BAP-Flag was IP’d and subject to proteomics analysis as a negative control. The Reactome FoamTree shows the over-represented functional pathways identified by proteomics ID of MIRO2-IP. **b**, Top 5 MIRO2 binders identified in the proteomics screen, according to spectral counts. **c**, PC3 cells transiently expressing BAP-Flag or MIRO2-Flag were subject to Flag-IP and analyzed by WB for co-IP of GCN1. A representative blot from n=3 experiments is shown. **d**, Endogenous MIRO2 or GCN1 were IP’d from PC3 cells, and analyzed for co-IP of GCN1 or MIRO2, respectively. IgG was used as an isotype control. A representative blot from n=3 experiments is shown. **e**, PLA assays to detect MIRO2/GCN1 proteins proximity in PC3 cells. A representative image (magnification 60X) is shown. Inset shows overlay between mitochondrially-targeted GFP and the PLA signal. **f**, WB of PC3 cells bearing MIRO2 (M2) or GCN1 (G1) knockdown. KO, knockout. **g**, PLA assays to detect MIRO2/GCN1 proteins proximity in PC3 cells. PLA using IgG and MIRO2 antibodies were used as negative controls. Specificity of the PLA was assayed by removing either MIRO2 (M2) or GCN1 (G1) and probing for the MIRO2/GCN1 interaction. PLA intensity per cell was represented as mean ± SEM (n=3). ***, p<0.001 by ANOVA and Dunnett’s post-test for multiple comparisons. **h**, GCN1 mRNA expression in benign prostate gland (n=20) or prostate adenocarcinoma (PRAD, n=69) in the Wallace dataset. ***, *p*=0.0004 by t-test with Welch’s correction. **j**, PLA assays to detect MIRO2/GCN1 proteins proximity in the indicated benign or cancer primary prostate epithelial cells. *Left*, PLA intensity per cell was represented as mean ± SD (n=9-14 independent fields). *Right*, Data are represented as mean ± SEM (n=3-5). *, p=0.0224 by unpaired t-test. **k**, The DepMap portal was searched for genetic cancer dependency on GCN1. Gene effect scores are derived from DEMETER2 or CERES.

We decided to follow up the top hit on the screen, General Control Non-derepressible (GCN1, Fig. 3b,c). Interestingly, GCN1 was included in both metabolism and cellular responses to extracellular stimuli pathways. In reciprocal co-IPs, GCN1 was bound to endogenous MIRO2 (Fig. 3d). The MIRO2-GCN1 interaction was detected by proximity ligations assays (PLA) in all prostate cancer cells models (Fig. 3e-g). As expected based on MIRO2 being anchored to the mitochondrial outer membrane, most of the PLA signal overlayed with a mitochondrially-targeted GFP (Fig. 3e, inset). Because both *MIRO2* and *GCN1* mRNA have higher expression in prostate cancer than normal adjacent prostate tissues (Fig. 1b and Fig. 3h), we compared the levels of MIRO2-GCN1 interaction across benign and cancer cell lines. We observed that the MIRO2-GCN1 interaction was nearly undetectable by PLA in diploid and immortalized benign prostate epithelial cells, compared to prostate cancer cell lines (Fig. 3i and Fig. S3d). Next, we wanted to show clinical relevance of this pathway using matched primary prostate epithelial cells isolated from patients. We found that the MIRO2-GCN1 interaction was exacerbated in primary prostate epithelial cancerous cells compared to benign cells (Fig. 3j and Fig. S3d).

GCN1 is the upstream activator of GCN2/EIF2AK4, a serine/threonine kinase that senses amino acid availability, redox status and actin dynamics cues in cells (29, 30). A recent study linked GCN1 to cell cycle regulation and proliferation in developing mice (31). However, there is no evidence of GCN1’s importance for cancer cells. Thus, as a first way to examine the role of GCN1 in controlling tumor cell growth, we mined the Cancer Dependency Map. Similar to MIRO2 (Fig. 1f), depletion of GCN1 in a large panel of pan-cancer cell lines by RNAi or CRISPR/Cas9 gene editing reduced cancer cell growth in most cell lines tested (Fig. 3k). As GCN1 ranked in the topmost dependent genes in > 90% of the cell lines, GCN1 is considered a common essential gene in cancer. Thus, GCN1 is upregulated in prostate cancer, is a critical pan-cancer mediator of cell growth and is a novel binding partner for MIRO2.

### GCN1 and GCN2 regulate prostate cancer cell growth

The importance of GCN2 was recently underscored by activation of GCN2 signaling in human breast, lung and liver tumors (32). Furthermore, GCN2 is essential for leukemia, melanoma and fibrosarcoma xenograft growth in mice (32–34). Interestingly, recent studies have linked mitochondrial stress to the activation of GCN2 and retrograde signaling to the nucleus in worms and mammalian cells (35, 36). Altogether, we postulate that GCN1 and GCN2 are key mediators of MIRO2’s effect on tumor cell growth in prostate cancer.

In order to determine if GCN1 and GCN2 control prostate cancer cell growth, we focused on a panel of prostate cancer cell lines from the Cancer Dependency Map. Similar to MIRO2 (Fig. 1g), GCN1 depletion via RNAi or CRISPR/Cas9 gene editing reduced cell growth (score < 0) in all cell lines tested (Fig. 4a). GCN2 depletion impaired cell growth as well, albeit to a lesser extent. To corroborate the results of the Cancer Dependency Map screen, we next generated stably depleted AR-independent prostate cancer cell lines (Fig. 4b). We found that stable GCN1 or GCN2 depletion reduced anchorage-dependent growth (Fig. 4c and Fig. S4a) and anchorage-independent cell growth (Fig. 4e). Furthermore, cell viability was lower in GCN1- or GCN2-depleted androgen-sensitive prostate cancer cell lines (Fig. S4b). Thus, we established that GCN1 and GCN2 are critical mediators of prostate cancer growth.

**Figure 4.**
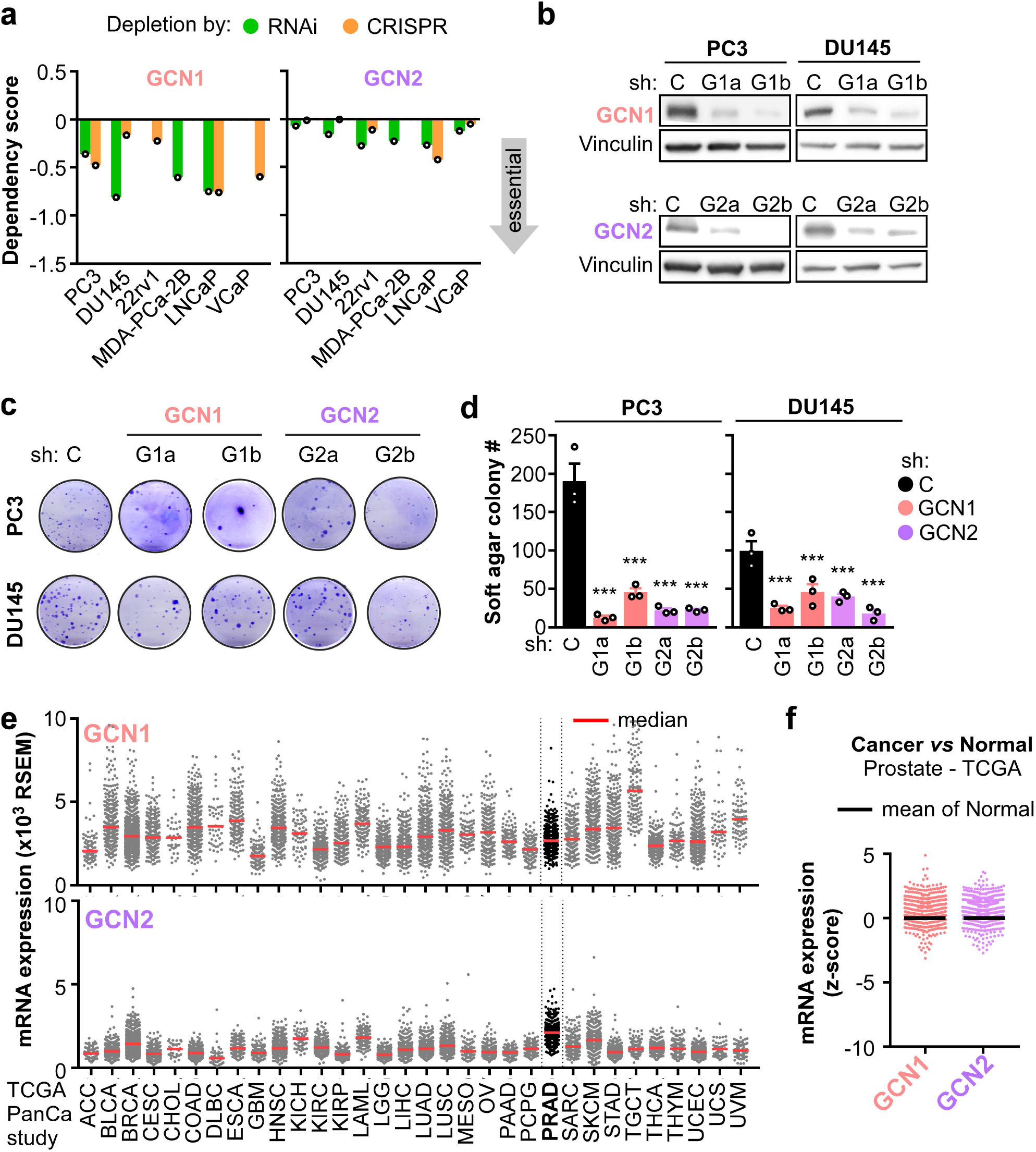
GCN1/2 regulation of prostate cancer cell growth. **a**, Dependency scores predicting the likelihood of GCN1 or GCN2 genes being essential on the prostate cancer cell lines indicated. Scores were retrieved from the Depmap portal (Broad Institute) from genome-wide RNAi or CRISPR based screens for vulnerabilities of cancer cells. **b-d**, Stable knockdown of GCN1 or GCN2 was achieved by shRNA in PC3 and DU145 cells. C, control sh; G1a/b, GCN1-targeting shRNAs (sequence a and b); G2a/b, GCN2-targeting shRNAs (sequence a and b). **b**, Representative blots showing the efficiency of knockdown are shown. **c**, Anchorage-independent growth in soft agar at 14 days post-plating. Colony # per well was quantitated and represented as mean ± SEM (n=3). ***, *p*<0.001 by ANOVA and Dunnett’s post-test for pairwise comparisons to control group. **d**, The TCGA database was interrogated for *GCN1/2* mRNA expression across tumor types. **e**, Relative expression of *GCN1/2* mRNA in cancer *versus* normal adjacent tissues on the prostate TCGA PanCancer study.

Next, we interrogated the TCGA PanCancer Atlas database at cBioPortal (18, 19) for expression of *GCN1* and *GCN2* in cancer. We found that *GCN1* and *GCN2* mRNA are expressed at variable levels across tumor types, with prostate adenocarcinoma (PRAD) showing one of the highest medians (Fig. 4e). Furthermore, higher *GCN1* and *GCN2* expression was present in cancer *versus* normal adjacent tissues in the TCGA and other patient cohorts (Fig. 4f and Fig. S4c). In terms of association between expression and patient prognosis, there was a trend towards higher *GCN1* expression being associated with poorer prognosis, but *GCN2* expression was not predictive of prognosis in prostate cancer (Fig. S4d). In summary, GCN1 and GCN2 are expressed across different tumor types and although highly expressed in prostate cancer, they do not predict patient prognosis.

### MIRO2 regulates GCN2 signaling

Given that loss of function of GCN1, GCN2 or MIRO2 impaired tumor cell growth, we next asked whether MIRO2 may modulate GCN1’s function as an activator of GCN2. GCN1 helps GCN2 sense deacylated tRNA that accumulate upon amino acid scarcity, leading to auto-phosphorylation of GCN2, and downstream phosphorylation of the eukaryotic translation initiation factor 2 alpha (eIF2α, Fig. 5a). Phosphorylated eIF2α shuts off global protein translation, while concomitantly inducing selective translation of the transcriptional activator 4 (ATF4) (29, 30). Due to the robust interaction between MIRO2 and GCN1, we hypothesized that MIRO2 regulates GCN2 signaling. When GCN2 activation was induced by amino acid starvation (AAS) we found that MIRO2 depletion impaired p-GCN2, p-eIF2α (Fig. 5b,c) and ATF4 levels (Fig. S5a) in prostate cancer. Thus, we have demonstrated for the first time that MIRO2 is a novel regulator of GCN2 signaling in prostate cancer cells.

**Figure 5.**
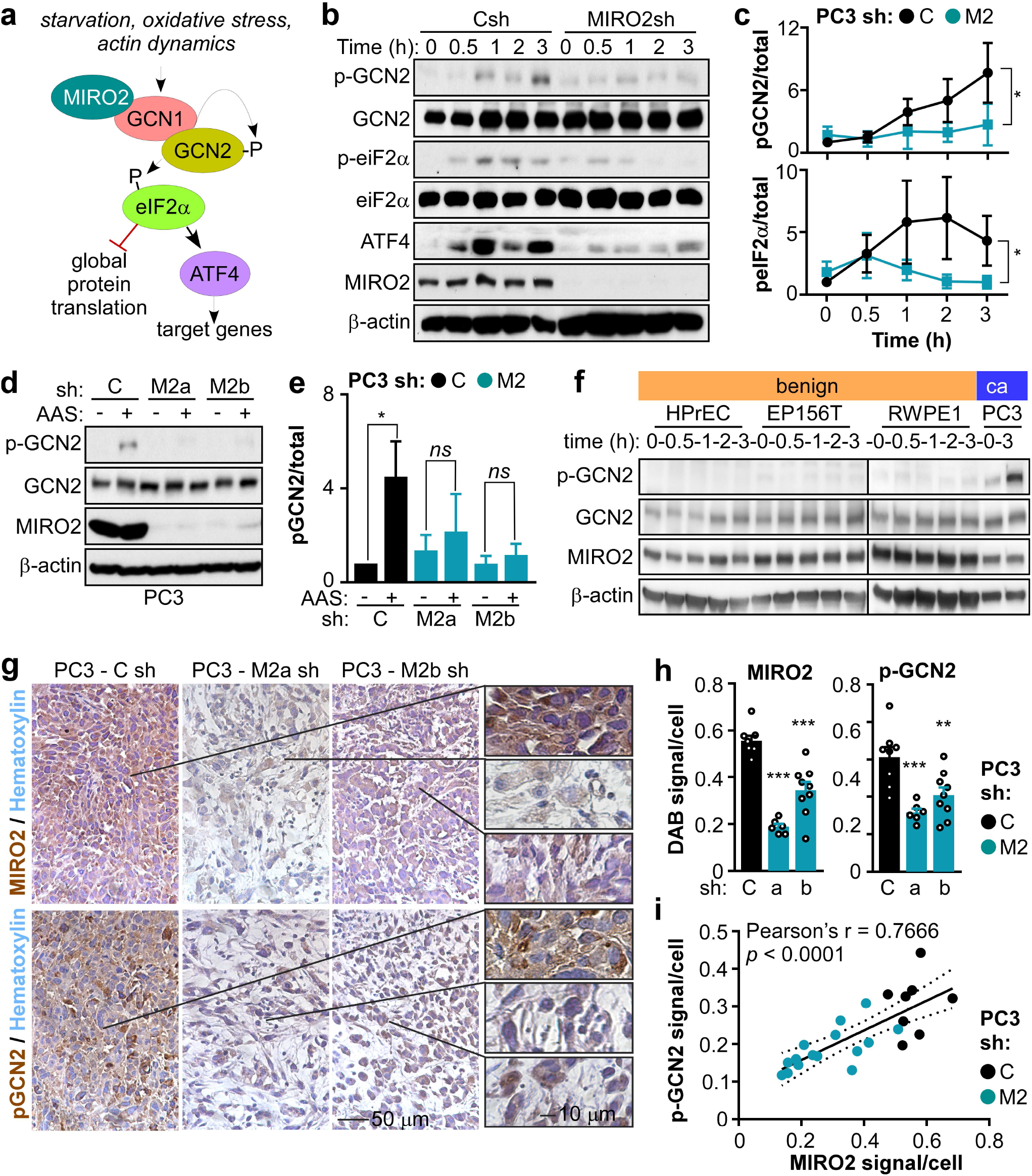
MIRO2 regulates GCN2 signaling. **a**, Canonical function of GCN1 as a co-activator of GCN2 signaling by amino acid/glucose starvation, oxidative stress, and actin dynamics. **b**, PC3 cells expressing Csh or MIRO2sh were incubated in amino acid starvation (AAS) conditions for 0-3 h and subject to WB. **c**, Densitometry analysis of the blots from b. Data is expressed as mean ± S.E.M. (n=3). *, *p*<0.05 by One-Way ANOVA and Tukey post-test. **d**, PC3 cells expressing Csh or MIRO2sh (two independent sequences, a and b) were incubated in full medium or amino acid starvation (AAS) conditions for 3 h and subject to WB. A representative blot of n=3 is shown. **e**, Densitometry analysis of the blots from d. Data is expressed as mean ± S.E.M. (n=3). *, *p*<0.05 by One-Way ANOVA and Dunnett’s post-test. **f**, Primary prostate epithelial (HPrEC), immortalized prostate epithelial (EP156T and RWPE1) or PC3 cells were incubated in full medium or amino acid starvation (AAS) conditions for 0-3 h and subject to WB. A representative gel is shown (n=3). **g**, Xenografts from PC3 cells expressing control (Csh) or MIRO2-sh (M2sh) were subject to IHC. Representative images at 10X magnification. **h**, DAB signal intensity per field was quantitated, normalized to the number of nuclei in the same field and represented as mean ± S.E.M. (n=3). ***, *p*=0.0001 by One-Way ANOVA and Dunnett’s post-test. **i**, Correlation between MIRO2 and p-GCN2 signal/cell in tumors from h.

Because the MIRO2-GCN1 interaction was undetectable in benign prostate epithelial cell lines, we next evaluated whether GCN2 could be differentially activated in prostate cancer cells. To this end, we compared the effects of AAS in human prostate epithelial cells of primary origin (HPrEC) or immortalized by telomerase bypass or viral transformation (EP156T and RWPE1). Indeed, we found that benign cell lines did not activate GCN2 upon amino acid starvation (Fig. 5f). These suggests that the MIRO2-GCN1-GCN2 signaling cascade is selectively activated in cancer cells over normal cells.

To test the relevance of this pathway *in vivo*, we examined the activation status of GCN2 in xenografts from control or MIRO2-depleted cancer cell lines. Phosphorylated-GCN2 levels were lower in MIRO2-depleted xenografts compared to control (Fig. 5g,h and Fig. S5c-f). Furthermore, p-GCN2 and MIRO2 levels were positively correlated (Fig. 5i). Overall, GCN2 is activated in prostate cancer xenografts in a MIRO2-dependent manner.

### MIRO2 controls the levels and activity of ATF4 in prostate cancer

In prostate cancer cells, we found that ATF4 induction upon AAS was dependent on MIRO2 expression (Fig. 4b and Fig. S5a). Interestingly, MIRO2 was required for maximal ATF4 induction in response to nutrient starvation both in AR-independent and androgen-sensitive cell lines (Fig. 6a,b). Transcriptional activity of an ATF4 luciferase reporter was dependent on MIRO2 expression (Fig. 6c). Importantly, ATF4 transcriptional activity was positively regulated by MIRO2 both in amino acid replete and amino acid depleted conditions (Fig. 6c,d). Overall, MIRO2 controls the levels and activity of ATF4 in AR-independent and androgen-sensitive prostate cancer cell lines.

**Figure 6.**
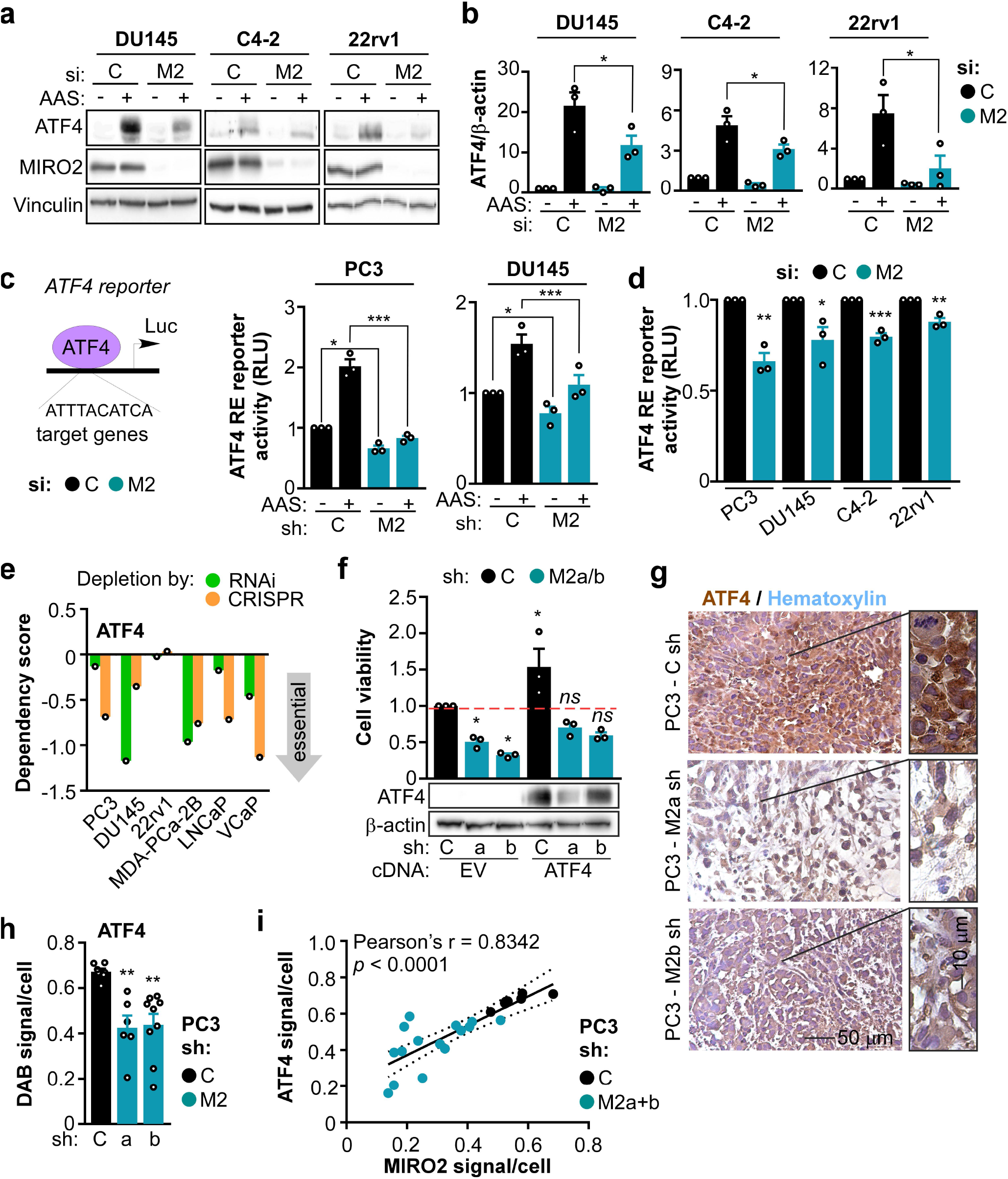
MIRO2 regulates ATF4 in prostate cancer. **a**, Cells were transfected with control (C) or MIRO2 (M2)-targeting RNAi, and 96 h later were and treated for 24 h in full or amino acid starved depleted medium (AAS). Representative WB are shown. **b**, Densitometry analysis of the blots from a. Data is expressed as mean ± S.E.M. (n=3). *, *p*<0.05 by One-Way ANOVA and Tukey post-test for multiple comparisons. **c**, The indicated cell lines expressing control (C) or MIRO2 (M2)-targeting RNAi were co-transfected with an ATF4-Firefly and a constitutive TK-Renilla reporter and treated for 24 h in full or amino acid starved depleted medium (AAS). *Left*, ATF4 reporter scheme. *Right*, ATF4 RE reporter activity was normalized to TK-Renilla and represented as mean ± SEM (n=3). *, *p*<0.01; ***, *p*<0.0001 by ANOVA and Tukey post-test for multiple comparisons. **d**, Cells were transfected as in (c) and basal levels of ATF4 reporter activity were measured in basal conditions (amino acids present). ATF4 RE reporter activity was normalized to TK-Renilla and represented as mean ± SEM (n=3). *, *p*<0.01; **, *p*<0.01; ***, *p*<0.0001 by unpaired t-test. **e**, Dependency scores predicting the likelihood of ATF4 being an essential gene on the prostate cancer cell lines indicated. Scores were retrieved from the Depmap portal (Broad Institute) from genome-wide RNAi or CRISPR based screens for vulnerabilities of cancer cells. **f**, PC3 cells expressing control (C) or MIRO2 (M2)-targeting shRNA were transfected with empty vector (−) or ATF4 cDNA (+) and assayed for cell viability 48 h later. Data are represented as mean ± SEM (n=3). *, p<0.01 by ANOVA and Dunnett’s post-test for multiple comparisons to the control group. *ns*, not significant. **g-i**, Xenografts from PC3 cells expressing control (Csh) or MIRO2-sh (M2sh) were subject to IHC. **g**, Representative images at 10X magnification. **h**, DAB signal intensity per field was quantitated, normalized to the number of nuclei in the same field and represented as mean ± S.E.M. (n=3). **, *p*<0.01 by One-Way ANOVA and Dunnett’s post-test. **i**, Correlation between MIRO2 and ATF4 signal/cell in tumors from h.

ATF4 controls the expression of a wide range of adaptive genes that allows cancer cells to adapt to excessive proliferation and demanding tumor microenvironment conditions (30). To determine whether ATF4 is required for prostate cancer cell growth, we mined the Cancer Dependency Map. Depletion of ATF4 via RNAi or CRISPR/Cas9 gene editing in a panel of prostate cancer cell lines reduced cell growth (score < 0) in all cell lines tested (Fig. 6e). As MIRO2 controlled ATF4 levels in prostate cancer, we next sought to assess whether lower levels of ATF4 in MIRO2-depleted cells could explain reduced prostate cancer cell growth. To this extent, we expressed ATF4 in control or MIRO2-depleted cells. We found that ATF4 expression increased tumor cell viability and partially rescued growth of MIRO2-depleted cells to control (Fig. 5f). Overall, MIRO2’s effect on regulating prostate cancer cell growth is partially mediated by ATF4.

To test the relevance of ATF4 *in vivo*, we examined the levels of ATF4 in xenografts from control or MIRO2-depleted cancer cell lines. ATF4 levels were lower in MIRO2-depleted xenografts compared to control (Fig. 6g,h). Furthermore, ATF4 and MIRO2 levels were positively correlated (Fig. 6i). Overall, ATF4 levels are dependent on MIRO2 expression in prostate cancer xenografts.

## DISCUSSION

In this study, we have shown that *MIRO2* mRNA expression is upregulated in prostate cancer and that higher levels of *MIRO2* correlate with poor patient survival in prostate cancer cohorts. This is in agreement with previous studies where *MIRO2* was overexpressed in cancer compared to benign tissues across disparate tumor types (13). Interestingly, the closely related *MIRO1* was down-regulated at the mRNA level, and did not correlate with patient survival. This is the first evidence that *MIRO1* and *MIRO2* are oppositely regulated in prostate cancer. Furthermore, we found that prostate cancer that recurred or progressed expressed higher levels of *MIRO2*, suggesting that *MIRO2* is important for prostate cancer tumor progression.

Our studies show that MIRO2 is required for growth of prostate cancer in cell lines representing AR-independent and androgen sensitive disease. Furthermore, MIRO2 depletion impaired tumor growth *in vivo*, which was associated with *in situ* decreased proliferation of tumor cells. Recently, MIRO2 was linked to regulation of cancer cell motility and metastasis (16). In these studies, Myc overexpression led to reprogramming of the mitochondrial network to fuel focal adhesion dynamics and motility. In contrast, we found that under basal conditions of endogenous expression of Myc, prostate cancer cells did not show alterations in gross mitochondrial distribution inside cells. One possibility is that Myc overexpression favors higher expression of MIRO1, MIRO2 and other mitochondrial trafficking proteins, thus enabling enhanced mitochondrial trafficking. Interestingly, our data indicates that MIRO1 is not required for regulating prostate cancer cell growth. To date, this is one of the first observations where MIRO1 and MIRO2 show divergent functions in mammalian cells.

Our proteomics approach identified novel protein binders of MIRO2 in prostate cancer cells. Previously known interactors of MIRO2 in neurons or fibroblasts included TRAK proteins that link MIRO2 to microtubules (37, 38). In our studies, MIRO2 did not co-precipitate with TRAK, nor did it alter mitochondrial distribution within cancer cells. Other protein binders for MIRO2 in non-tumorigenic cells include the E3 ubiquitin ligase Parkinson Protein 2 (PARK2) and the kinesin-like protein motor Centromere Protein F (CENPF) (26, 39). Of note, association to PARK2 occurs exclusively under severe stress induced by treatment with mitochondrial uncouplers (26), and association to CENPF is exclusively detected in M phase of the cell cycle (39). The fact that these previously known MIRO binding partners were not present in our IP screen may be because our studies were carried out in prostate cancer cells that were not stressed or arrested on M phase of the cell cycle. Alternatively, cancer cells may have a largely non-overlapping interactome with normal cells. This idea warrants further exploration, as there are no other available proteomic studies identifying MIRO2 binding partners in normal *versus* cancer cells.

In this study, we focused our efforts in characterizing the function of GCN1, a previously unknown protein binder of MIRO2. Our data shows that GCN1 is needed for anchorage-dependent and – independent growth of AR-independent and androgen-sensitive prostate cancer cells. Furthermore, GCN1 was a common essential gene in a large panel of pan-cancer cell lines. Altogether, these are the first evidence implicating GCN1 in controlling cancer cell biology. Interestingly, GCN1 was also overexpressed in prostate cancer compared to benign tissue and the interaction of MIRO2-GCN1 was increased in prostate cancer *versus* immortalized prostate epithelial cells. Thus, our data suggests that targeting the MIRO2-GCN1 axis may selectively affect tumors while minimizing side effects on normal tissues.

The canonical function of GCN1 is to activate the kinase GCN2 in response to amino acid starvation, accumulation of globular actin or oxidative stress (29). However, recent evidence points to GCN2-independent effects of GCN1 in the control of inflammatory and tissue remodeling responses (40). In our studies, we found that MIRO2 was required for activation of GCN2 signaling in prostate cancer cells. In agreement with previous studies in fibrosarcoma (32), GCN2 was activated in prostate cancer xenograft tumors. Such activation of GCN2 in tumors may result from: (i) a challenging microenvironment depleted of oxygen and nutrients, or (ii) tumor metabolic rewiring to redirect available nutrients into anabolic pathways to support tumor proliferation and biomass expansion. In prostate cancer xenograft tumors, MIRO2 levels were correlated with activation of GCN2, reinforcing that MIRO2 is a novel regulator of GCN2 signaling in prostate cancer. Furthermore, GCN1 and GCN2 were required for anchorage-dependent and –independent growth of prostate cancer cells. Thus, GCN1’s canonical function as an activator of GCN2 is important for prostate cancer cell growth.

An open question that stems from our studies is what are the upstream signals that trigger MIRO2-dependent GCN1/2 activation in prostate cancer cells? As MIRO2 is a mitochondrial protein located in the outer mitochondrial membrane, it may be ideally located to sense and transduce mitochondrial fitness signals back to the nucleus by engaging GCN1/2-dependent retrograde signaling. In support of this idea, studies in worms and non-transformed mammalian cells have shown that ROS or amino acid depletion engage GCN2 retrograde signaling (35, 36). We do not know whether this mechanism is at play in prostate cancer cells, or how local pools of mitochondrially-produced ROS and amino acids levels engage MIRO2-GCN1. However, the fact that the MIRO2-GCN1 interaction, and MIRO2-dependent activation of GCN2 selectively occur in cancer cells suggests that transformed cells benefit from having a pool of GCN1/2 locally activated at the mitochondrial surface.

In terms of the downstream mechanisms that mediate tumor cell growth downstream of MIRO2, GCN1 and GCN2, we show that ATF4 partially mediates MIRO2 effects in prostate cancer cell growth. Interestingly, MIRO2 controls ATF4 transcriptional activity both in amino acid replete and amino acid depleted conditions. In prostate cancer cells, ATF4 is essential for cell growth, and ATF4 expression partially rescues cell growth of MIRO2-depleted cells. Furthermore, ATF4 levels in prostate cancer xenografts are correlated to MIRO2 expression. Overall, this indicates that MIRO2 positively regulates ATF4 levels and that contributes to prostate cancer cell growth. While ATF4 is the best characterized downstream protein induced by GCN2 activation, our studies cannot rule out a potential role of MIRO2 in controlling additional stress sensing kinases that converge on eIF2α, including the ER sensor EIF2AK3 (41, 42). Because MIRO2 has been linked to ER-mito signaling in normal cells (24), the possibility that MIRO2 is a broad regulator of kinases converging on eIF2α/ATF4 will need further exploration.

In summary, we identified a novel signaling pathway centered on MIRO2 that is required for growth of prostate cancer cells *in vitro* and *in vivo*. MIRO2 relies on activation of GCN1/2 signaling and ATF4 induction to support prostate cancer cell growth. *MIRO2* and *GCN1* are both overexpressed in prostate cancer patient cohorts, and the MIRO2-GCN1 interaction occurs selectively in prostate cancer cells. We therefore propose that targeting the MIRO2-GCN1 axis may be a valuable strategy to halt prostate cancer growth. This novel strategy would have application to both AR-independent and androgen-sensitive tumors.

## Supporting information

Supplemental Fig S1-S5, Table S1, Supp Methods and references

## ACKNOWLEDGEMENTS

The authors acknowledge Monika Dzieciatkowska for proteomics sample preparation and analysis, Dana Davis at the Flow Cytometry Core for cell cycle analyses, and the Histology Core for histology services. The authors thank Dr. Andrew Thorburn for critically reading the manuscript.

