## Supplemental Fig S1-S5, Table S1, Supp Methods and references for "MIRO2 regulates prostate cancer cell growth via GCN1-dependent stress signaling"

Running title: MIRO2 and GCN1 regulate prostate cancer growth

Keywords: MIRO2, GCN1, mitochondria, prostate cancer

Contents:

Supplemental Figures (Figures S1-S5)

Supplemental Tables (Tables S1-S3; Tables S2, S3 are supplied as separate Excel files)

Supplemental Methods

Supplemental References

**SUPPLEMENTAL FIGURES**

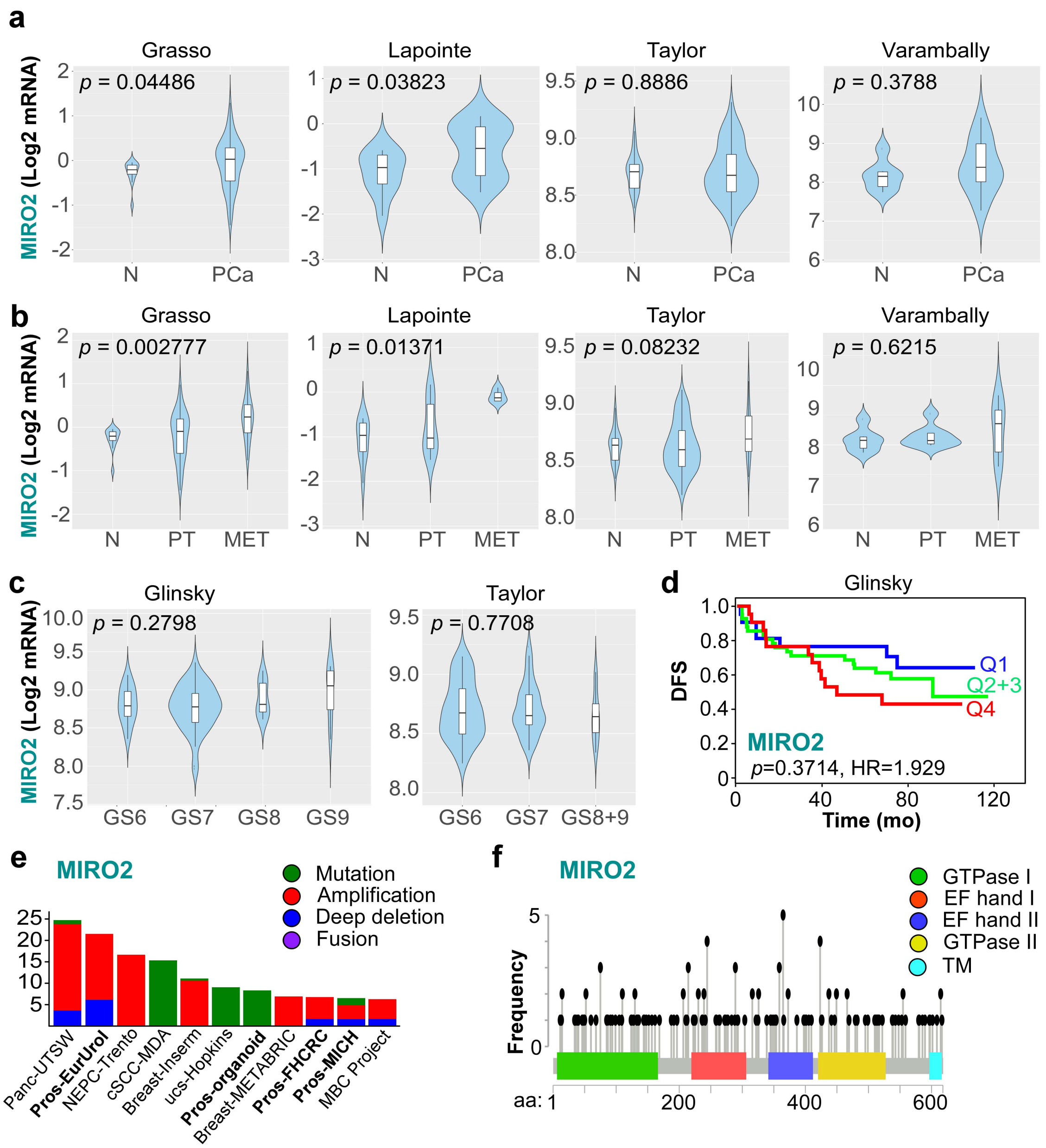


Figure S1, related to Figure 1. **MIRO1/2 alterations in cancer**. **a-c**, Violin plots depicting *MIRO2* mRNA expression in prostate cancer (PCa) *versus* non-tumoral (N) specimens (a), according to progression status (b) or according to Gleason grade (c) were generated using CANCERTOOL. N, non-tumoral; PCa, prostate cancer; PT, primary tumor; MET, metastatic prostate cancer; GS, Gleason score. Means were compared compared by t-test (panel a) or ANOVA (panels b,c). **d**, Kaplan-Meier analyses based on *MIRO2* mRNA expression in human primary tumor samples. The Glinsky dataset was split at quartiles, where Q4>Q3>Q2>Q1. Q1 vs Q4 curves were compared with a Mantel-Cox test. Hazard ratio (HR) was calculated with a Cox proportional hazards regression model. DFS, disease-free survival. **e**, The cBioPortal database was interrogated for MIRO2 copy number and mutation status across all available studies (curated set of non-redundant studies, n=48,091 patients). Studies where >5% of cases were altered are shown. **f**, Mutation frequency for *MIRO2* was obtained from studies in (e).

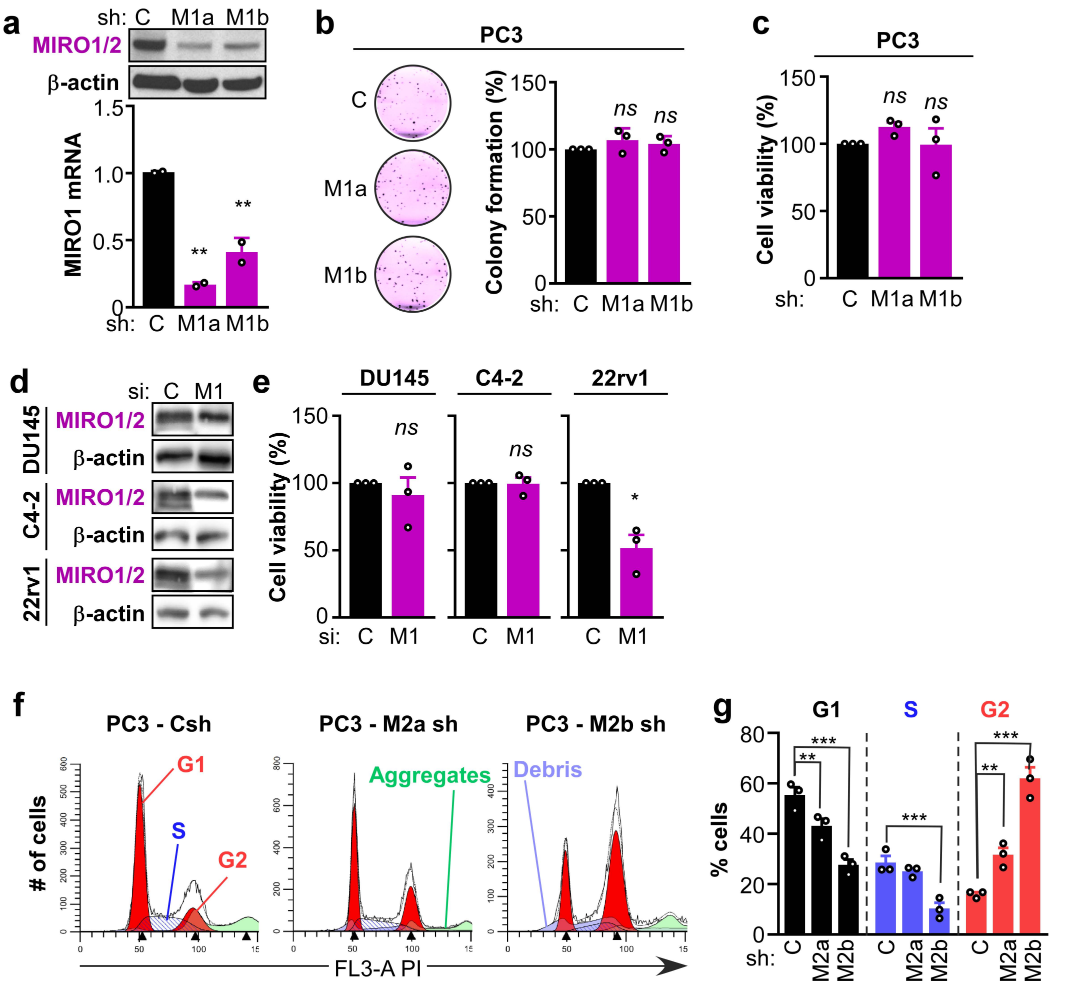


Figure S2, related to Figure 2. **MIRO1 depletion does not affect tumor cell intrinsic phenotypes**. **a-c**, Stable knockdown of MIRO1 was achieved by shRNA in PC3 cells. C, control sh; M1a/b, MIRO1-targeting shRNAs (two independent sequences, a and b). **a**, Representative blots showing the efficiency of knockdown using a MIRO1/2 antibody are show (top). Relative mRNA levels of MIRO1 were assayed by qPCR and represented as mean ± SEM (n=3). **, p=0.0041-0.0015 by One-Way ANOVA and Dunnett's post-test for multiple comparisons. **b**, Anchorage-dependent growth at 14 days post-plating. *Left*, Representative scans of stained colonies. *Right*, quantitation of colony # per well, relative to control (C) and represented as mean ± SEM (n=3). *ns*, not significant by One-Way ANOVA and Dunnett's post-test for multiple comparisons. **c**, Cell viability was determined by a MT-Glo assay and relativized to the control (Csh). Data is represented as mean ± SEM (n=3). *ns*, not significant by One-Way ANOVA and Dunnett's post-test for multiple comparisons. **d-e**, Cells were transiently transfected with control (C) or MIRO1 (M1)-targeting siRNA and cell viability was determined 6 days later by a MT-Glo assay and relativized to the control (Csi). **d**, representative WB showing the level of depletion of MIRO1. **e**, Data is represented as mean ± SEM (n=3). *, $p$=0.0392 by unpaired t-test with Welch's correction. *ns*, not significant. **f**, Cells were stained with propidium iodide and analyzed for DNA content by flow cytometry. Representative cell cycle profiles are shown. **g**, Data from (f) is represented as mean ± SEM (n=3). **, $p$<0.01; ***, $p$<0.001 by One-Way ANOVA and Dunnett's post-test for multiple comparisons.

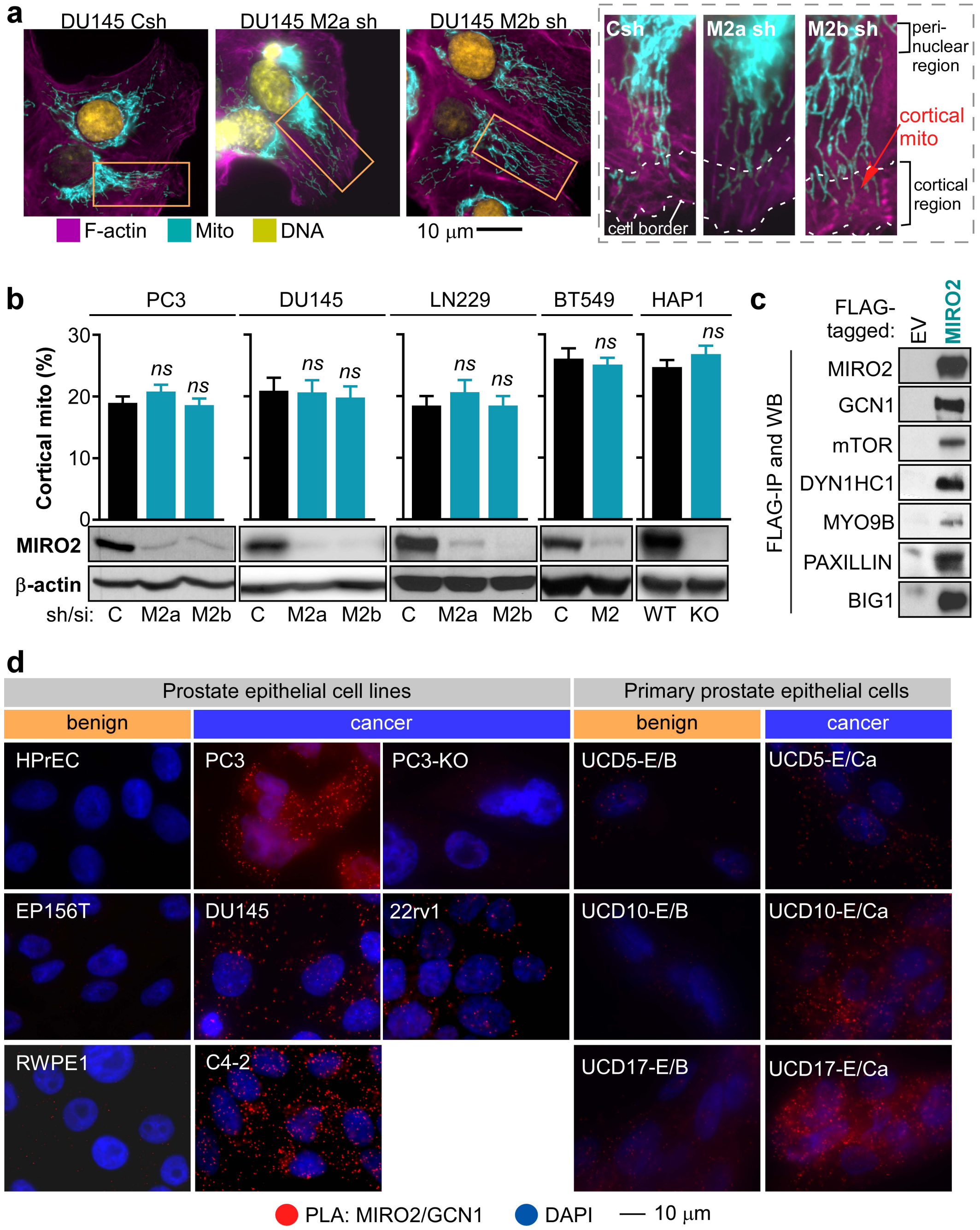


Figure S3, related to Figure 3. **MIRO2 regulates mitochondrial dynamics in stress conditions. a,** The indicated cell lines were fixed and immunostained with markers for filamentous actin (F-actin), mitochondria (Mito) and DNA and analyzed under the microscope. Right, zoomed panels show mitochondria in cortical and peri-nuclear regions. **b,** The indicated cell lines were control (C) or MIRO2 knockdown was achieved transiently (si) or stably (sh, 2 independent sequences, M2a and M2b) growing exponentially were analyzed for cortical mitochondria. Data represent cortical mitochondria signal normalized to total mitochondria mass per single cell (mean ± S.E.M., n>100). All comparisons were non-significant by ANOVA (p>0.05). **c,** PC3 cell lysates containing FLAG-empty vector (EV) or FLAG-MIRO2 were subject to IP with FLAG antibodies and analyzed for co-IP of the indicated proteins by Western blotting. Only IP fractions are shown. **d,** Representative images for MIRO2/GCN1 interaction by PLA. *Left,* Benign prostate epithelial (HPrEC, EP156T and RWPE1) or prostate cancer (PC3 and DU145) cells were compared. *Right,* Matched primary patient cells isolated from benign (E/B) or carcinoma (E/Ca) areas from the same patient were compared.

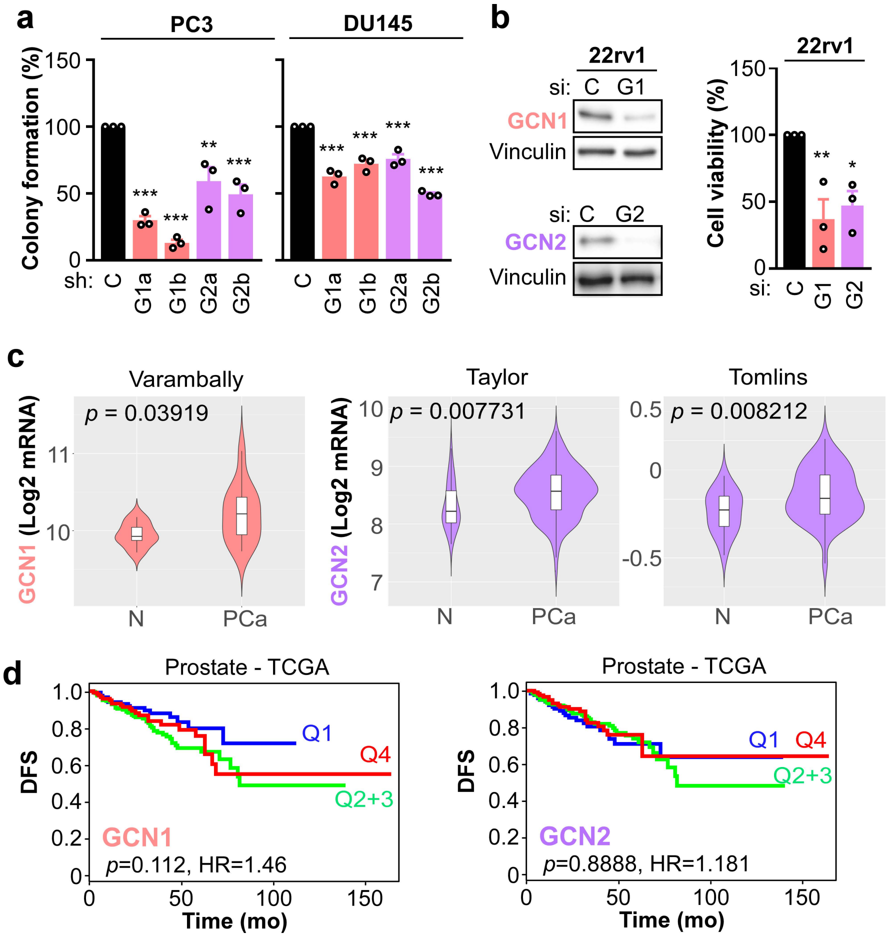


Figure S4, related to Figure 4. **GCN1/2 regulate mPC cell growth**. **a**, Stable knockdown of GCN1 or GCN2 was achieved by shRNA in PC3 and DU145 cells. C, control sh; G1a/b, GCN1-targeting shRNAs (sequence a and b); G2a/b, GCN2-targeting shRNAs (sequence a and b). Anchorage-dependent growth at 14 days post-plating. Quantitation of colony # per well relative to control, represented as mean ± SEM (n=3). ***, $p<0.001$ by ANOVA and Dunnett's post-test for pairwise comparisons against to group. **b**, Cells were transiently transfected with control (C), GCN1 (G1)- or GCN2 (G2)-targeting siRNA and cell viability was determined 6 days later by a MT-Glo assay and relativized to the control (Csi). *Left*, representative WB showing the level of depletion of MIRO2. *Right*, data is represented as mean ± SEM (n=3). *, p = 0.0219; **, p = 0.01 by One Way ANOVA and Dunnett's post test for multiple comparisons against control. **c**, Violin plots depicting *GCN1/2* mRNA expression in prostate cancer (PCa) *versus* non-tumoral (N) specimens were generated using CANCERTOOL. Datasets with statistically significant comparisons are shown. Means were compared compared by t-test. **d**, Kaplan-Meier analyses based on *GCN1/2* mRNA expression in human primary tumor samples. The TCGA dataset was split at quartiles, where Q4>Q3>Q2>Q1. Q1 vs Q4 curves were compared with a Mantel-Cox test. Hazard ratio (HR) was calculated with a Cox proportional hazards regression model. DFS, disease-free survival.

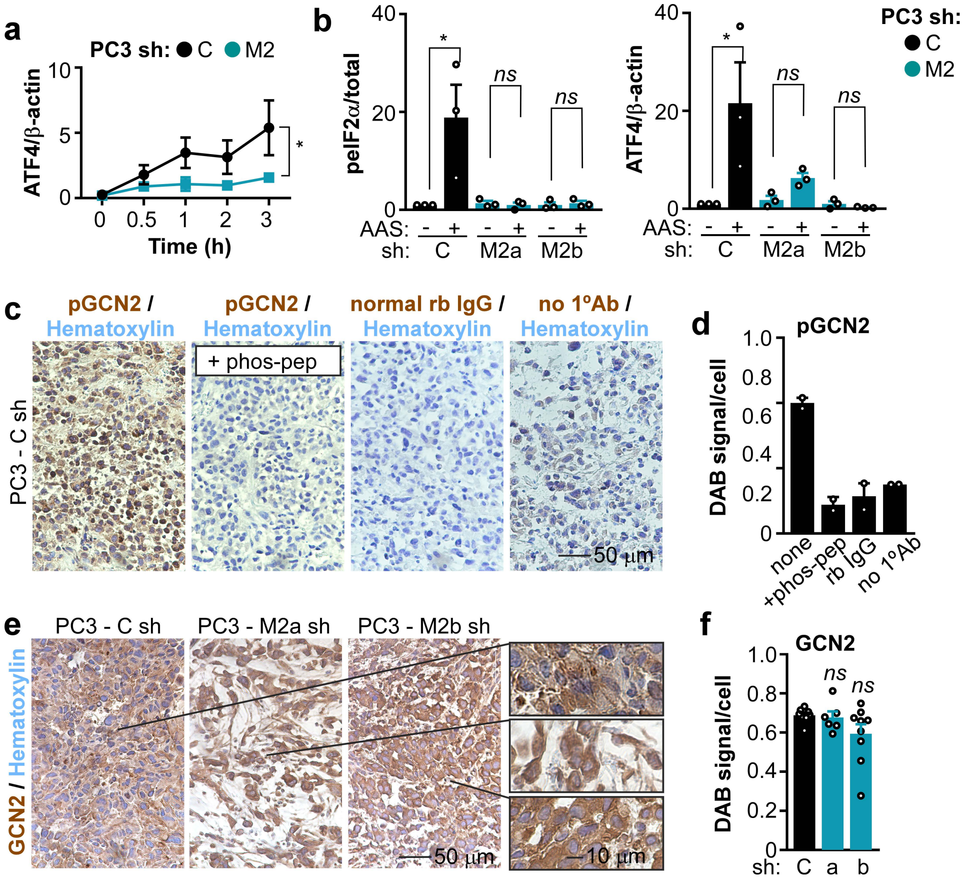


Figure S5, related to Figure 5. **MIRO2 regulates GCN2 signaling**. **a**, PC3 cells expressing Csh or MIRO2sh were incubated in amino acid starvation (AAS) conditions for 0-3 h and subject to WB. Densitometry analysis of the blots is expressed as mean ± S.E.M. (n=3). *, $p<0.05$ by One-Way ANOVA and Tukey post-test. **b**, PC3 cells expressing Csh or MIRO2sh (two independent sequences, a and b) were incubated in full medium or amino acid starvation (AAS) conditions for 3 h and subject to WB. Densitometry analysis is expressed as mean ± S.E.M. (n=3). *, $p<0.05$ by One-Way ANOVA and Dunnett's post-test. **c,** Xenografts from PC3 cells expressing control (Csh) were subject to IHC. IgG was used as an isotype control. To demonstrate specificity of the p-GCN2 antibody, the antibody was blocked with phospho T899-GCN2 peptides (p-pep, 200X molar excess for 1 h) previous to IHC. Representative images at 10X magnification. **d**, DAB signal intensity per field was quantitated, normalized to the number of nuclei in the same field and represented as mean ± S.E.M. (n=2). **e,** Xenografts from PC3 cells expressing control (Csh) or MIRO2-targeting shRNA (M2sh) were subject to IHC. Representative images at 40X magnification (insets at 100X magnification). **f**, DAB signal intensity per field was quantitated, normalized to the number of nuclei in the same field and represented as mean ± S.E.M. (n=3). *ns*, not significant by One-Way ANOVA and Dunnett's post-test for comparisons against control group.

### SUPPLEMENTAL TABLES

Table S1. **Top 50 putative binders of MIRO2.** Putative interactors of MIRO2 were identified by Proteomics ID of a FLAG-MIRO2 IP. Common contaminants found in the Crapome database were removed. Proteins that were undetected (nd) in the FLAG-BAP control were ranked based on average spectral counts on the MIRO2-IP. These top 50 putative binders of MIRO2 are represented below. The row containing GCN1's data is highlighted. Rep 1-3, replicates 1-3. Asterisks (*) indicate hits that were independently validated by FLAG and Western blot detection using endogenous antibodies against that target (see Fig. S3c). *nd*, undetected in FLAG-BAP control.

| | | **FLAG-MIRO2** | | | FLAG-MIRO2 |
|---|---|---|---|---|---|
| ***Accession Number*** | ***FLAG-BAP*** | ***rep 1*** | ***rep 2*** | ***rep 3*** | ***mean*** |
| MIRO2_HUMAN | *nd* | 767 | 406 | 672 | 615 |
| GCN1_HUMAN * | *nd* | 251 | 235 | 235 | 240.3333333 |
| RN213_HUMAN | *nd* | 175 | 172 | 178 | 175 |
| HUWE1_HUMAN | *nd* | 163 | 148 | 154 | 155 |
| XPO1_HUMAN | *nd* | 124 | 40 | 115 | 93 |
| UBR4_HUMAN | *nd* | 90 | 69 | 79 | 79.33333333 |
| BAG6_HUMAN | *nd* | 88 | 68 | 77 | 77.66666667 |
| MDN1_HUMAN | *nd* | 85 | 70 | 75 | 76.66666667 |
| XPO2_HUMAN | *nd* | 93 | 31 | 80 | 68 |
| USP9X_HUMAN | *nd* | 72 | 43 | 53 | 56 |
| MTOR_HUMAN * | *nd* | 64 | 53 | 44 | 53.66666667 |
| SYFA_HUMAN | *nd* | 57 | 46 | 55 | 52.66666667 |
| TBA1C_HUMAN | *nd* | 54 | 47 | 55 | 52 |
| NU205_HUMAN | *nd* | 60 | 51 | 43 | 51.33333333 |
| TECR_HUMAN | *nd* | 63 | 43 | 47 | 51 |
| IPO5_HUMAN | *nd* | 56 | 26 | 64 | 48.66666667 |
| CMC1_HUMAN | *nd* | 53 | 30 | 57 | 46.66666667 |
| ECM29_HUMAN | *nd* | 58 | 41 | 26 | 41.66666667 |
| SFXN3_HUMAN | *nd* | 45 | 45 | 35 | 41.66666667 |
| BIG1_HUMAN * | *nd* | 49 | 38 | 31 | 39.33333333 |
| IPO7_HUMAN | *nd* | 51 | 15 | 52 | 39.33333333 |
| QCR2_HUMAN | *nd* | 42 | 31 | 44 | 39 |
| RPN2_HUMAN | *nd* | 39 | 39 | 38 | 38.66666667 |
| NDUS1_HUMAN | *nd* | 44 | 24 | 45 | 37.66666667 |
| CMC2_HUMAN | *nd* | 47 | 17 | 47 | 37 |
| MON2_HUMAN | *nd* | 39 | 38 | 34 | 37 |
| BIRC6_HUMAN | *nd* | 44 | 30 | 36 | 36.66666667 |
| DNJA1_HUMAN | *nd* | 33 | 35 | 41 | 36.33333333 |
| EHD4_HUMAN | *nd* | 38 | 28 | 41 | 35.66666667 |
| GET4_HUMAN | *nd* | 36 | 39 | 30 | 35 |
| ATM_HUMAN | *nd* | 46 | 29 | 29 | 34.66666667 |
| FADS2_HUMAN | *nd* | 36 | 32 | 35 | 34.33333333 |

| | | | | | |
|---|---|---|---|---|---|
| EHD1_HUMAN | *nd* | 39 | 27 | 36 | 34 |
| SQOR_HUMAN | *nd* | 32 | 33 | 35 | 33.33333333 |
| ACSL3_HUMAN | *nd* | 43 | 17 | 39 | 33 |
| AT1A1_HUMAN | *nd* | 37 | 24 | 33 | 31.33333333 |
| MCM7_HUMAN | *nd* | 34 | 14 | 44 | 30.66666667 |
| FAF2_HUMAN | *nd* | 32 | 28 | 30 | 30 |
| CPT1A_HUMAN | *nd* | 35 | 21 | 32 | 29.33333333 |
| M2OM_HUMAN | *nd* | 34 | 26 | 28 | 29.33333333 |
| CO1A2_HUMAN | *nd* | 21 | 32 | 32 | 28.33333333 |
| COPB_HUMAN | *nd* | 39 | 12 | 34 | 28.33333333 |
| DHB12_HUMAN | *nd* | 33 | 25 | 25 | 27.66666667 |
| XPOT_HUMAN | *nd* | 36 | 16 | 29 | 27 |
| GBF1_HUMAN | *nd* | 29 | 25 | 26 | 26.66666667 |
| LTN1_HUMAN | *nd* | 37 | 27 | 16 | 26.66666667 |
| MYO9B_HUMAN * | *nd* | 21 | 30 | 27 | 26 |
| SFXN1_HUMAN | *nd* | 31 | 27 | 20 | 26 |
| KNTC1_HUMAN | *nd* | 30 | 25 | 22 | 25.66666667 |
| IPO4_HUMAN | *nd* | 37 | 9 | 31 | 25.66666667 |

Table S2. **Comprehensive list of proteins identified in MIRO2-IP.** Putative interactors of MIRO2 were identified by Proteomics ID of a FLAG-MIRO2 IP. Common contaminants found in the Crapome database were removed. Spectral counts are provided for a single FLAG-BAP control and 3 replicate MIRO2-IP samples in the “raw” tab. Proteins that were undetected in the FLAG-BAP control were ranked based on average spectral counts on the MIRO2-IP and are presented in the “top binders” tab. The row containing GCN1’s data is highlighted. Rep, replicate.

*Table S2 is provided as a separate Excel file.*

Table S3. **Pathways over-represented in MIRO2-IP.** Putative interactors of MIRO2 were identified by Proteomics ID of a FLAG-MIRO2 IP. Proteins present with more than 10 spectral counts in the MIRO2-IP were analyzed for over-representation pathway analysis using the Reactome database. Pathways are ranked by number of entities identified in the MIRO2-IP. The total number of entities, with associated *p* values and FDR are provided.

*Table S3 is provided as a separate Excel file.*

### SUPPLEMENTAL METHODS

**Cell culture.** Human prostate adenocarcinoma (C4-2, DU145, and PC3), glioblastoma (LN229), breast ductal carcinoma (BT549), and prostate epithelial cells (HPrEC, EP156T, RWPE1) were obtained from ATCC, and maintained in culture according to the supplier's specifications. Parental and MIRO2 KO near-haploid chronic myelogenous leukemia (HAP1) cells derived from the KBM-7 cell line were obtained from Horizon Discovery Group (Cat#HZGHC89941). Both WT and MIRO2 KO cell lines were sequenced to confirm the CRISPR/Cas9 editing of the MIRO2 allele, which had a 13 bp deletion in exon 6 that led to a stop codon at amino acid 51. Master stocks of cell lines were authenticated using small tandem repeat (STR) analysis (The University of Colorado Cancer Center Cell Culture core) and tested for mycoplasma with the ATCC Universal Mycoplasma Detection Kit (ATCC# 301012K). Cells were cultured for a maximum of 8 weeks (or 8 passages for prostate epithelial cells) and mycoplasma testing was repeated upon freezing stable shRNA cell lines and prior to injecting in mice.

**Antibodies and reagents.** Antibodies to MIRO2 (Thermo Fisher # PA552960, diluted 1:2000), GCN1 (Abcam # ab86139, diluted 1:1,000), FLAG (clone M-2, Sigma-Aldrich # F3165, diluted 1:2000), Total GCN2 (CST #65981, diluted 1:2000), Phospho-GCN2 T899 (Abcam ab75836 diluted 1:1000), Phospho-eIF2α S51 (Cell Signaling Technologies #3398S diluted 1:1000), Total-eIF2α (clone D7D3, CST #5324S diluted 1:3000), ATF4 (clone D4B8, CST # 11815, diluted 1:1000), mTOR (clone#7C10, CST # 2983, diluted 1:1000), DYN1HC1 (Protein Tech # 12345-1-AP, diluted 1:5000), MYO9B (Thermo Fisher # PA5-18524, diluted 1:250), PAXILLIN (clone 5H11, Thermo Fisher # AHO0492, diluted 1:1000), BIG1 Abcam # ab72061, diluted 1:1000), Vinculin (clone E1E9V, CST #13901, diluted 1:400,000), and β-actin (clone AC-15, Sigma-Aldrich # A5441, diluted 1:200,000) were used for Western blotting. Secondary antibodies for Western blotting were from CST and highly cross-adsorbed secondary antibodies for immunofluorescence were from Thermo Fisher.

Amino Acid Starved (AAS) Media contained 10% dialyzed FBS (Gemini Bio-Products #100-108) in RPMI 1640 Media Medium w/o Amino Acids, Sodium Phosphate (US Biologicals # R8999-04A) supplemented with 1 g of sodium bicarbonate.

**Plasmids and transfections.** TrueORFGold pCMV6-Entry-myc-FLAG plasmids encoding MIRO2 were from Origene (#RC204823). The ATF4 reporter (ATF4-RE-FireflyLuc) was a kind gift of Dr. Winklhofer, and was characterized previously (1). Mouse ATF4 was a gift from David Ron (Addgene plasmid # 21845). Cells were transfected with 1 μg pDNA plus 2 μl X-treme gene HP (Sigma #06366236001) for 24 hours in complete medium, washed, and subjected to the indicated treatments.

**Gene silencing.** For transient knockdown experiments by siRNA, pooled siRNA nucleotides were transfected at 10-15nM concentrations in the presence of RNAiMAX Lipofectamine (Thermo Fisher) at a 1:2.5 siRNA:Lipofectamine ratio. The following RNAi Dharmacon SMARTpools were used: control, non-targeting siRNA SMARTpool (D-001810-10), *MIRO1* (L-010365-01), *MIRO2* (L-008340-01), *GCN1* (L-027130-01) or *GCN2* (L-005314-00). After 72-96 hours, cells were validated for target protein knockdown by Western blotting and processed for subsequent experiments.

For stable knockdown experiments in human cells, two independent shRNA (acquired from the University of Colorado Cancer Center Functional Genomics Core) were used: *MIRO1* sequence a (TRCN 0000353760), *MIRO1* sequence a TRCN 0000353761; *MIRO2* sequence a (TRCN 0000072919), *MIRO2* sequence b (TRCN 0000072921); *GCN1* sequence a (TRCN 0000336602), *GCN1* sequence b (TRCN 0000336604); *GCN2* sequence a (TRCN 0000304216), *GCN2* sequence b (TRCN 0000300850). An empty pLKO-based lentivirus was used as control. PC3 and DU145 cells stably expressing shRNA or empty pLKO, were generated by infection with lentiviral particles, followed by 2 weeks of selection in the presence of puromycin at 2μg/mL. Cells bearing stable knockdowns of GCN1 and GCN2 were selected and maintained with puromycin at 2 µg/ml, 50 µM ß-Mercaptoethanol, and 1X MEM Non-essential Amino Acids (Thermo Fisher #11140-050).

**Western blotting.** Protein lysates were prepared in 25 mM Tris HCl pH 7.5, 100 mM NaCl, 5 mM EDTA, 10% glycerol, 1% Triton X-100 containing EDTA-free Protease inhibitor cocktail (Thermo Fisher # 78438). For all experiments analyzing phospho-proteins, phosphatase inhibitor tablets were used (Thermo Fisher # 32957). Lysates were sonicated and precleared by centrifugation at 17,000 x*g* for 10 min at 4°C. Protein concentrations were determined via Pierce 660nm Protein Assay (Thermo # 22660). Equal amounts of protein lysates were separated by SDS gel electrophoresis, transferred to polyvinylidene difluoride membranes, blocked in 5% non fat milk (CST # 9999) diluted in TBST buffer (20 mM Tris HCl, pH 7.5, 150 mM NaCl, 0.1% Tween-20), and further incubated with primary antibodies diluted in 5% BSA/TBST overnight at 4 °C. After washing in TBST, membranes were incubated with HRP-conjugated secondary antibodies (1:1000 dilution in 5% BSA/TBST) for 1 h at room temperature. For membranes from endogenous and FLAG co-IP experiments, HRP-conjugated anti-rabbit IgG confirmation specific (1:1000 dilution in 5% BSA/TBST) antibodies were used. Membranes were washed with TBST and proteins were visualized via chemiluminescence in a KwikQuant Imager system (Kindle Biosciences LLC) using Ultra Digital-ECL Substrate Solutions (Kindle Biosciences # R1100). Intensity of bands relative to a control band was quantitated with the KwikQuant Image Analyzer 3.1 software (Kindle Kindle Biosciences LLC).

**Endogenous Co-IP.** Protein lysates were prepared in IP buffer (25 mM Tris-HCl pH 7.5, 100 mM NaCl, 5 mM EDTA, 10% glycerol, 1% Triton X-100 containing EDTA-free Protease inhibitor cocktail (Thermo Fisher # 78438) by rotating for 1 h at 4°C. Lysates were precleared by centrifugation at 17,000 x*g* for 10 min at 4°C and protein concentrations were determined via Pierce 660nm Protein Assay (Thermo # 22660). Five hundred µg of protein rotated overnight at 4°C with either 5 µg of anti-GCN1 (Abcam # ab86139), 1 µg anti-MIRO2 (Thermo Fisher # PA552960), or normal Rabbit IgG (CST # 2729) control matched to the highest antibody concentration. Protein A agarose beads (CST # 9863, 20µL per sample) were equilibrated in IP buffer and added to protein-antibody mixture and rotated at 4°C for 1 h. Beads were washed 4 times in IP buffer and protein was eluted by boiling in 1X Laemmli buffer (50 mM Tris-HCl pH 6.8, 10% glycerol, 2% SDS, 0.005% bromophenol blue, 2% β-mercaptoethanol). Samples were then analyzed via Western blotting as described above.

**FLAG IP.** $4 \times 10^6$ cells were plated in 15-cm culture dishes in growth medium without antibiotics and allowed to adhere overnight at 37°C in a 5% $CO_2$ incubator. Transfection complexes containing 18µg of pDNA (FLAG-MIRO2 or FLAG-empty vector), 32 µL of X-tremeGENE HP DNA transfection reagent (Roche # 6366546001) and 3.2 mL Opti-MEM were incubated for 30 min at room temperature and added to cells. Protein lysates were prepared in IP buffer (25 mM Tris-HCl (pH 7.5), 100 mM NaCl, 5 mM EDTA, 10% glycerol, 1% Triton X-100 containing EDTA-free Protease inhibitor cocktail) by rotating for 1 h at 4°C. Lysates were precleared by centrifugation at 17,000 x*g* for 10 min at 4°C and protein concentrations were determined via Pierce 660nm Protein Assay. Anti-FLAG M2 beads (Sigma-Aldrich # A2220, 20µL per sample) were equilibrated in IP buffer and added to 500 µg of protein lysates. For FLAG-BAP samples, 500 ng of FLAG-BAP fusion protein (Sigma-Aldrich # P7582) was added to the protein-bead mixture. Protein-bead mixtures were rotated for 2 h at 4°C. Beads were washed 4 times in IP buffer and protein was eluted in 1X Laemmli buffer (50 mM Tris-HCl pH 6.8, 10% glycerol, 2% SDS, 0.005% bromophenol blue, 2% β-mercaptoethanol). Samples were then analyzed via Western blotting as described above, or subject to proteomics ID (see below).

**Proteomics ID.** The samples were loaded onto a 1.5 mm thick NuPAGE Bis-Tris 4−12% gradient gel (Thermo Fisher) and visualized with SimplyBlue™ SafeStain (Thermo Fisher) stain. Each lane of the gel was divided into 6 equal-sized bands, and proteins in the gel were digested as follows. The pieces were destained in 200 µL of 25 mM ammonium bicarbonate in 50 % v/v acetonitrile for 15 min and washed with 200 µL of 50% (v/v) acetonitrile. Disulfide bonds were reduced by dithiothreitol, and cysteine residues were alkylated with iodoacetamide. Modified sequencing grade trypsin (Promega) was added 1:100 w/w and the samples were digested overnight at 37 C. The digestion was stopped by addition of 5% formic acid. Peptides were extracted two times from the gel plugs using 1% formic acid in 50% acetonitrile. The collected extractions were pooled with the initial digestion supernatant, desalted, and concentrated on Thermo Scientific Pierce C18 Tip. Samples were analyzed on an Orbitrap Fusion mass spectrometer (Thermo Fisher Scientific) coupled to an Easy-nLC 1200 system (Thermo Fisher Scientific) through a nanoelectrospray ion source. Peptides were separated on a self-made C18 analytical column (100 µm internal diameter, x 20 cm length) packed with 2.7 µm Cortecs particles. After equilibration with 3 µL 5% acetonitrile 0.1% formic acid, the peptides were separated by a 70 min a linear gradient from 6% to 38% acetonitrile with 0.1% formic acid at 400nL/min. LC mobile phase solvents and sample dilutions used 0.1% formic acid in water (Buffer A) and 0.1% formic acid in 80 % acetonitrile (Buffer B) (Optima™ LC/MS, Fisher Scientific, Pittsburgh, PA). Data acquisition was performed using the instrument supplied Xcalibur™ (version 4.1) software. Survey scans covering the mass range of 350–1800 were performed in the Orbitrap by scanning from m/z 300-1800 with a resolution of 120,000 (at m/z 200), an S-Lens RF Level of 30%, a maximum injection time of 50 milliseconds, and an automatic gain control (AGC) target value of 4e5. For MS2 scan triggering, monoisotopic precursor selection was enabled, charge state filtering was limited to 2–7, an intensity threshold of 2e4 was employed, and dynamic exclusion of previously selected masses was enabled for 45 seconds with a tolerance of 10 ppm. MS2 scans were acquired in the Orbitrap mode with a maximum injection time of 35 ms, quadrupole isolation, an isolation window of 1.6 m/z, HCD collision energy of 30%, and an AGC target value of 5e4.

MS/MS spectra were extracted from raw data files and converted into mgf files using a Proteome Discoverer Software (ver. 2.1.0.62). These .mgf files were then independently searched against human database using an in-house Mascot server (Version 2.6, Matrix Science). Mass tolerances were +/- 10 ppm for MS peaks, and +/- 25 ppm for MS/MS fragment ions. Trypsin specificity was used allowing for 1 missed cleavage. Met oxidation, protein N-terminal acetylation, and peptide N-terminal pyroglutamic acid formation were allowed as variable modifications. Scaffold (version 4.8.4, Proteome Software, Portland, OR, USA) was used to validate MS/MS based peptide and protein identifications. Peptide identifications were accepted if they could be established at greater than 95.0% probability as specified by the Peptide Prophet algorithm. Protein identifications were accepted if they could be established at greater than 99.0% probability and contained at least two identified unique peptides.

**Reactome pathway analysis.** The comprehensive list proteins identified as putative interactors of MIRO2 by Proteomics ID (section above) were used as starting point. First, common background contaminants found in proteomics samples (e.g. keratins) were identified via Crapome at http://www.crapome.org/ (2) and removed from the dataset. Second, counts in the control (FLAG-BAP) IP were substracted from counts for each of the 3 replicates for FLAG-MIRO2 IPs (Table S2). Then the average for the background substracted MIRO2 IPs was calculated, and any proteins with averages less than 10 spectral counts were excluded. The resulting list of protein identifiers was submitted for overrepresentation pathway analysis against Reactome version 73 on 19/07/2020, using the Reactome database (https://reactome.org/). Probability scores were corrected for false discovery rate by the Benjamini-Hochberg method. Table S3 presents the results for overrepresentation analysis.

**mRNA quantitation.** mRNA levels for human MIRO1 were determined by qPCR. Briefly, RNA was extracted with a PureLink RNA Mini Kit (ThermoFisher) following the in-column DNA digestion protocol. One µg of RNA was reversed transcribed using random hexamer primers and a High Capacity cDNA Reverse Transcription Kit (ThermoFisher #4368814). Two µl of cDNA diluted 1:5 were used as template for qPCR reaction with TaqMan Gene Expression assays. Pre-designed Taqman assays were human MIRO1 (Hs00430253_m1) and eukaryotic 18S rRNA (4352930E). For relative quantitation, the ΔΔCt method was used.

**Analyses of expression and cancer cell dependency from public databases.** The TCGA PanCancer studies containing tumor expression data for *MIRO1, MIRO2, GCN1* and *GCN2* mRNA (RNA-seq values) were downloaded from the cBioPortal for Cancer Genomics (http://www.cbioportal.org/) (3, 4) and plotted with GraphPad Prism 7.0 software. Genomic alterations for *MIRO1* and *MIRO2* were obtained from all available studies in cBioPortal (n=48,091 patients) and plots displaying the frequency of alterations and lollipop mutations were downloaded from cBioPortal.

For prostate cancer-specific analyses, the TCGA PanCancer prostate adenocarcinoma dataset was accessed via the cBioPortal and interrogated for expression of particular genes in cancer *vs* normal adjacent tissues, or according to disease free status. Violin plots for association between mRNA expression and tumor progression or Gleason grade, and Kaplan-Meier survival curves were generated using the CANCERTOOL platform at http://genomics.cicbiogune.es/CANCERTOOL/index.html (5).

For analyses of cancer cell dependency on a particular gene, we searched the Cancer Dependency Map website from Broad Institute. CERES or DEMETER2 gene effect scores from DepMap public release 19Q3 were downloaded from https://depmap.org/portal/download/ (Oct 10 2019) (6). Data were plotted using GraphPad Prism 7.0.

**Cell growth assays.** For anchorage-dependent growth, 200 cells were plated in triplicate on 6-well plates and incubated for 10 days (DU145) or 14 days (PC3) in a humidified incubator set to 37°C and 5% $CO_2$. Fresh medium was added every 2-3 days. The cells were stained with a 0.5% w/v crystal violet/methanol solution for 30 minutes at 23°C. Plates were scanned in a EPSON Perfection V19 Model J371A scanner and images were imported into FIJI software to manually count the colonies per well using the Multipoint selection tool.

For anchorage-independent growth, 3000 cells were plated in 0.5 ml of 0.35% agar/medium on top of 0.5 ml of a solid layer of 0.5% agar/medium and allowed to solidify for 20 min at room temperature. Medium was added to the top and replaced every 3 days. Cells were allowed to grow for 14 days in a humidified incubator set to 37°C and 5% $CO_2$. At the end, live cells were stained with a 2 mg/ml solution of MTS for 2 hs in a humidified incubator set to 37°C and 5% $CO_2$. Wells were inspected by bright field microscopy under an Olympus CKX41 inverted microscope with a UpPlanFL N 4X/0.13 PhP objective, and live colonies (brownish) in all focal planes were counted.

For cell viability, $5 \times 10^5$ cells (DU145, 22rv1, and C4-2) were transfected with 15nM of pDNA (Non-targeting control, GCN1, or GCN2) with 7.5 µL Lipofectamine RNAi/Max (Themofisher # 13778075) in 1mL of Opti-MEM. The cells were plated in 6-well plates and allowed to adhere for 48 hours at 37°C in a 5% CO2 incubator. Transfected cells (3,000/well) were plated in triplicate on 96-well plates in 200 µL of media. The cells were incubated for 72 hours at 37°C in a 5% CO2 incubator. Changes in cell viability were determined using the RealTime-Glo™ MT Cell Viability Assay (Promega, Madison, WI) using a FilterMax F5 Multi-Mode Microplate reader (Molecular Devices, San Jose, CA).

**Cell cycle analyses.** PC3 cells were seeded in triplicate at 125,000 cells/well in 6-well plates and grown at 37°C in a 5% CO2 incubator for 48 h. Cells were collected via trypsinization, washed one time with PBS, and the pellet was resuspended in 500µL of PBS. Cells were added drop by drop to ice cold 70% ethanol while vortexing at max speed. Samples were stored at -20°C until day of Flow cytometry analysis. On the day of Flow cytometry analysis, fixed cells were collected via centrifugation and washed one time in PBS. Cells were resuspended in 250µL of DNA staining solution containing 20µg/mL Prodidium Iodide (Thermo Fisher # P3566), 200µg/mL RNAse A (NEB # T3018L), 0.1% Triton X-100 (Fisher # BP151-500) in PBS. Cells were incubated at room temperature for 30 min protected from light, and then placed on ice until analysis via flow cytometry. Cells suspensions were analyzed on Beckman Coulter Gallios using a 488nm laser line. Voltage was set to 314 and approximately 10,000 events were collected per sample. Percent of cells in each cell cycle phase was calculated using a ModFit LT analysis.

**Proximity ligation assays.** Cells were seed at 40,000-75,000 cells/well on 18mm sterile glass coverslips in 12-well plates. Cells were allowed to adhere to coverslips at 37°C in a 5% CO2 incubator for 72 hours. Cells were fixed in Formalde-Fresh (Fisher # SF93-4) for 15 min at room temperature. Cells were washed 3 times with PBS and permeabilized with 0.1% Triton X-100 (Fisher # BP151-500)/PBS for 5 minutes at room temperature. Cells were washed 3 times in PBS and blocked in 5% normal goat serum (Gibco # 16-210-064)/0.3 M Glycine (Fisher # BP381-500)/PBS for 1 h at room temperature. Coverslips were transferred to slides with parafilm and PLA blocking buffer was added to coverslips for 1 h at room temperature. Coverslips were transferred back to 12-well plate and incubated in primary antibody overnight at 4°C. Normal IgG were used as isotype controls and were diluted to achieve the same concentration as the diluted mouse anti-MIRO2 or rabbit anti-GCN1 antibodies respectively. Antibodies used were: anti-GCN1 (Abcam # ab86139, diluted 1:1000, final concentration 0.2 µg/ml), anti-MIRO2 (Abnova # H00089941-A01, diluted 1:1000, final concentration 0.1 µg/ml), normal rabbit IgG (CST # 2729, diluted 1:5000, final concentration 0.2 µg/ml to match anti-GCN1), mouse IgG1 Isotype Control (CST # 5415, diluted 1:10000, final concentration 0.1 µg/ml to match anti-MIRO2). Coverslips were then transferred back to slides with parafilm and placed in a humidified chamber. The rest of the PLA procedure was followed according to manufacturer's protocol, and at the end of the procedure slides were washed and mounted in 4,6-diamidino-2-phenylindole (DAPI)-containing Prolong Gold mounting medium (ThermoFisher).

Images were taken on an EVOS XL Core Imaging System at 60X magnification, using an Olympus PlanApo N 60X/1.42 Oil objective, and EVOS FL LED cubes. Texas Red FL LED cube has Ex=585/29 nm and Em=624/40 nm and DAPI FL LED cube has Ex=357/44 nm and Em=447/60 nm. For each sample at least 10 random fields containing 100 cells were imaged. Images were imported into FIJI software, deconvoluted into red and blue channels. The red channel was used to quantitate the red fluorescence intensity using the Analyze/Measure feature in FIJI. A small empty region on the red channel field which did not contain visible signal was used to measure the background and extrapolate it to the full area of the field. The blue channel was used to manually count the nuclei per fiel. PLA signal of a field was obtained by substracting the red background signal from the red fluorescence signal of each field. A normalized PLA Signal/Cell value was calculated by dividing the PLA signal by the number of cells in the same field.

**Immunofluorescence and cortical mitochondria quantitation.** Cells under the various conditions tested were fixed in formalin/PBS (4% final concentration) for 15 min at 22 °C. Cells were then permeabilized with 0.1% Triton X-100/PBS for 5 min, washed, and incubated in 5% normal goat serum (NGS, Vector Labs) diluted in 0.3 M glycine/PBS for 60 min. Primary antibodies against mitochondria (clone MTC02, Thermo Fisher # MA512017, diluted 1:500) were added in 5% NGS/0.3 M glycine/PBS and incubated for 18 h at 4 °C. After three washes in PBS, secondary antibodies anti-mouse conjugated to Alexa594 were diluted 1:500 in 5% NGS/0.3 M glycine/PBS and added to cells for 1 h at 22 °C. Where indicated, F-actin was stained with phalloidin Alexa488 (Thermo Fisher # A12379, diluted 1:250) for 30 min at 22 °C. Slides were washed and mounted in 4,6-diamidino-2-phenylindole (DAPI)-containing Prolong Gold mounting medium (Thermo Fisher #P36931).

At least seven random fields were analyzed by fluorescence microscopy in an EVOS XL Core Imaging System at 60X magnification, using an Olympus PlanApo N 60X/1.42 Oil objective, and EVOS FL LED cubes. Texas Red FL LED cube has Ex=585/29 nm and Em=624/40 nm, YFP FL LED cube has Ex=500/24 nm and Em=524/27 nm and DAPI FL LED cube has Ex=357/44 nm and Em=447/60 nm. Mitochondria/F-actin composite images were analysed in ImageJ as described (7). Briefly, the F-actin channel was used to manually label the cell boundary and a belt extending from the boundary towards the inside of the cell was marked as the 'cortical mask'. This cortical mask was subsequently applied to the mitochondria channel to measure intensity at the cortical region, and normalized to total mitochondria intensity per cell and cell area. A minimum of 30 cells per experiment were analyzed and data from 3 independent experiments (total 100 cells) were pooled to obtain mean values and perform statistical testing.

**Immunohistochemistry.** At the end of the experiment, animals were euthanized by $CO_2$ inhalation followed by cervical dislocation, and tumors were removed by necropsy. Tumors were rinsed in PBS and a 2 mm thick slice perpendicular to the largest diameter and spanning the central area of the tumor was dissected and fixed in neutral formalin (Fisher Scientific # SF93-4). Tumor slices were immediately given to UCD Research Histology Shared Resource (University of Colorado, Aurora, CO, USA) to be processed for paraffin embedded and cut into five-micrometer serial sections. Slides were warmed at 50°C for 20 minutes; deparaffinized in xylene for 20 minutes, then xylene/ethanol 1:1 for 5 minutes; and rehydrated in alcohol series (100%, 95%, 90%, 70%, 50%, 30% ethanol and $dH_2O$, 5 minutes each). Antigen retrieval was done in citrate-based solution at pH 6.0 (Vector Laboratories # H-3300) or Tris-based solution at pH 9.0 (Vector Laboratories # H-3301) or in a pressure cooker for 5 minutes, followed by cooling to room temperature. Next, slides were washed once in PBS or TBST for 5 minutes and incubated in Bloxall (Vector Laboratories #30150) for 10 minutes. Slides were then washed 3 times with PBS or TBST (5 minutes each) and blocked in 2.5% normal horse serum (Vector Laboratories #30022) for 20 minutes at room temperature. Primary antibody was diluted in SignalStain Antibody Diluent (Cell Signaling Technologies #8112, for Ki-67 antibody) or 2% BSA in TBST (phospho-antibodies) or PBS (total antibodies). Antibodies used were: Ki-67 (Cell Signaling Technologies #9027, diluted 1:400), total-GCN2 (Abcam # ab134053, diluted 1:100), Phospho-Thr899-GCN2 Affinity Biologicals # AF8154, diluted 1:50), MIRO2 (Proteintech #11237-1-AP, diluted 1:1000), ATF4 (Proteintech #10835-1-AP, diluted 1:200). Slides were incubated in a humidified chamber for 2 hours at room temperature with the antibody dilutions and then washed 3 times in wash buffer (TBST for phospho-antibodies, PBS for total antibodies) for 5 minutes each, incubated with IMPRESS Horse anti-Rabbit polymer (Vector Laboratories #30026) at room temperature for 30 minutes, and washed 3 times for 5 minutes each. Slides were developed with a DAB+ substrate chromogen system (Vector Laboratories # SK-4103) for 1-30 minutes, rinsed in $dH_2O$, and stained with Hematoxylin QS (Vector Laboratories # H-3404) for 5 seconds. Slides were dehydrated in $dH_2O$, 30%, 50%, 70%, 90%, 95%, and 100% ethanol (5 minutes each); immersed in xylene for 15 minutes; and mounted with VectaMount Permanent mounting medium (Vector Laboratories # H-5000).

Semi-quantitative determination of protein expression using DAB staining was carried out as described in Ref. (8). Briefly, DAB stained sections were imaged in a Nikon 50i upright microscope, DXM1200F camera, and ACT-1 software (Nikon). For each field, images were imported into FIJI (Fiji is Just Image J) software for Windows and deconvoluted using the Image/Color/Color Deconvolution plugin for hematoxylin and DAB (H DAB). The DAB window was thresholded (minimum: 0, maximum: based on the average or the control group, see below), and analyzed for mean grey value and area. The H window was used to manually count the nuclei with the Multipoint function. The DAB signal/cell was calculated by dividing the DAB mean grey value obtained by the number of nuclei (positive for hematoxylin) on the same field. The maximum threshold value was empirically determined for each antibody stain by obtaining the best threshold on the DAB window for each control field, and averaging it. This average maximum threshold was 205 for MIRO2, 190 for p-GCN2 and total GCN2, and 165 for ATF4.

For Ki67 quantification, 5 representative images were taken for each xenograft. The percent of Ki67-positive cells per image was determined by dividing the number of Ki67-positive nuclei by the total number of nuclei (positive for hematoxylin) in the same field, and multiplying by 100.

Specificity of the phosho-GCN2 antibody was determined by blocking with phospho-GCN2 (Thr899) blocking peptides (Affinity Biologicals # AF8154-BP). Blocking peptides were resuspended at 10 mg/ml in $dH_2O$ and incubated at a 200X molar excess with an aliquot of the 1° antibody AF8154 for 1 h at room temperature. A control reaction had $dH_2O$ (same volume as the peptide tube) with an aliquot of the 1° antibody AF8154 for 1 h at room temperature. Samples were centrifuged at maximum speed in a refrigerated centrifuge for 15 min to pellet immune complexes, and the supernatant was transferred to a new tube and used for IHC of tissues.
